# The Effects of DMSO on DNA Conformations and Mechanics

**DOI:** 10.1101/2025.03.31.646418

**Authors:** Koen R. Storm, Caroline Körösy, Enrico Skoruppa, Stefanie D. Pritzl, Pauline J. Kolbeck, Willem Vanderlinden, Helmut Schiessel, Jan Lipfert

## Abstract

Dimethyl sulfoxide (DMSO) is a polar aprotic solvent used in a wide range of applications, including uses as a drug and in drug delivery, as a solvent for fluorescence dyes, and in enzymatic reactions that process DNA. Consequently, many assays contain low concentrations (≤ 10%) of DMSO. While it is well known that DMSO lowers the melting temperature of DNA, its effects on DNA conformations and mechanical properties below the melting temperature are unclear. Here we use complementary single-molecule techniques to probe DNA in the presence of 0-60% DMSO. Magnetic tweezers force-extension measurements find that the bending persistence length of DNA decreases moderately and linearly with DMSO concentrations up to 20 vol%, by (0.43 ± 0.02)% per %-DMSO, respectively. Magnetic tweezers twist measurements demonstrate a reduction in melting torque in the presence of DMSO and find that the helical twist of DNA remains largely unchanged up to 20% DMSO, while even higher concentrations slightly unwind the helix. Using AFM imaging, we find a moderate compaction of DNA conformations by DMSO and observe a systematic decrease of the mean squared end-to-end distance by 1.2% per %-DMSO. We use coarse grained Monte Carlo simulations of DNA as a semi-flexible polymer with a variable density of flexible segments or bubbles, representing DMSO-induced local defects or melting, to rationalize the observed behavior. The model quantitates the effects of introducing locally flexible regions into DNA and gives trends in line with the magnetic tweezers and AFM imaging experiments. Our results show that addition of up to 50% DMSO has a gradual effect on DNA structure and mechanics and that for low concentrations (≤ 20%) the induced changes are relatively minor. Our work provides a baseline to understand and model the effects of DMSO on DNA in a range of biophysical and biochemical assays.

**STATEMENT OF SIGNIFICANCE:** Dimethyl sulfoxide (DMSO) is a widely used polar aprotic solvent. It is employed e.g. as a drug and in drug delivery, as a solvent for small molecules, and as an additive to enzymatic reactions that process DNA. Despite its wide-spread use, its effects on DNA conformations and properties are not well understood and often neglected. We use single-molecule manipulation with magnetic tweezers and AFM imaging to probe DNA in the presence of 0-60% DMSO. We find that DMSO increases the flexibility of DNA, leading to more compact conformations. Up to 50% DMSO, the induced changes are gradual and approximately linear. Our results provide a quantitative baseline to understand, model, and optimize assays with DNA in the presence of DMSO.

## INTRODUCTION

Dimethyl sulfoxide (DMSO) is a polar aprotic solvent and widely used in a large range of applications (1). It is used clinically to treat interstitial cystitis (2–4) and acts as a topical analgesic, anti-inflammatory, and antioxidant (5,6). DMSO is frequently used as a solvent for stock solutions of e.g. small molecule drugs and fluorescent dyes. As a result, many biological and biochemical assays contain DMSO in low concentration as an inadvertent component (7,8). In addition, DMSO is employed as a cryo- and radioprotectant (9–11), and frequently used as an additive to increase yield in biochemical reactions, in particular in PCR and topoisomerization assays (12–14). DMSO is well known to lower the melting temperature of DNA (15,16), however, it is less clear how it affects overall DNA structure and mechanical properties (17–19). While the effects of low concentrations of DMSO in aqueous buffers on DNA mechanics are often neglected, some studies suggest that they can induce significant changes of DNA stiffness. For example, recent work using AFM imaging reported a > 4-fold reduction in DNA bending stiffness in 3% DMSO (20).

Here we use single-molecule assays to directly probe the effects of DMSO on the overall conformations, bending stiffness, and helical twist of DNA. In a first set of measurements, we use magnetic tweezers (MT) to determine the stretching and twist response of DNA in the presence of varying concentrations of DMSO, ranging from 0% to 60% DMSO (v/v) in a background of (near-) physiological PBS buffer. In MT, DNA molecules are tethered between a flow cell surface and ∼1 µm diameter superparamagnetic beads (21,22). External magnets enable the application of precisely calibrated stretching forces (23) and control of the DNA linking number (21,24). From force-extension measurements, we find a moderate and initially linear decrease of the bending persistence length with increasing DMSO concentration, by (0.43 ± 0.02)% per %-DMSO. From extension-rotation measurements at constant applied force, we observe essentially no change in DNA twist, except for the highest DMSO concentrations, but a reduction of the melting torque upon addition of DMSO.

In a second set of experiments, we use AFM imaging of linear DNA in air to directly observe the conformations of free dsDNA in the presence of varying concentrations of DMSO in the absence of applied forces and torques (25–28). We observe a moderate compaction of DNA upon addition of DMSO and a corresponding average reduction of the end-to-end distance, by 1.2% per %-DMSO.

To rationalize the observations from our single molecule measurements in a quantitative model, we perform extensive coarse-grained Monte Carlo simulations of double-stranded DNA, both with applied external forces and in free solution. While our coarse-grained simulations with implicit solvent do not treat individual bases and do not provide direct insights into the interactions of DMSO with DNA bases, they have the advantage to enable simulations of DNA molecules in the length range used experimentally (∼ kbp) (29–35). We model the effect of DMSO by adding flexible defect segments or “bubbles”, i.e. DNA positions that are much more flexible than the regular segments and thus can accommodate local bending more readily than standard B-form DNA. We find that an increasing density of bubbles leads to a decrease of the apparent persistence length and end-to-end lengths, consistent with the experimental observations for DNA in the presence of DMSO and in agreement with the notion that the presence of kinks renormalizes the bending persistence length (36,37).

## MATERIALS AND METHODS

### Chemicals and preparation of dilution series

DMSO was purchased from Sigma-Aldrich (Dimethyl sulfoxide, for molecular biology (≥ 99.9%); catalogue number D8418, Sigma-Aldrich). The DMSO solutions were prepared in 1× phosphate buffered saline (PBS, consisting of 10 mM phosphate buffer, pH 7.4, with 137 mM NaCl and 2.7 mM KCl; catalogue number P3813, Sigma-Aldrich) for concentrations of 1-60% (v/v) DMSO. Refractive indices of the solutions were determined using an Abbe refractometer (Abbe Refractometer 3T, Atago; at *λ* = 589.3 nm and 21 °C) for MT analysis (Supplementary Figure S1).

### DNA construct and flow cell preparation for magnetic tweezers measurements

For MT measurements, we used a functionalized 20.6 kbp DNA construct that was described previously (38). The DNA construct has (≈600 bp) PCR-generated handles with multiple biotin or digoxigenin groups ligated to the ends of the unmodified central sequence of the DNA. Streptavidin-coated, 1.05 µm diameter superparamagnetic beads (MyOne, Thermo Fisher Scientific) are attached to the DNA constructs by binding to the biotin moieties, by adding 1 μL of picomolar DNA stock solution to 2 μL of the magnetic beads in 200 μL 1× PBS.

Flow cells are prepared using two microscope coverslips (24 × 60 mm; Carl Roth). The bottom coverslip was functionalized with (3-Glycidoxypropyl)trimethoxysilane and incubated with 100 μL of a 5000× diluted stock solution of 1 µm diameter polystyrene beads (Polysciences) in ethanol, to be used as reference beads. Alternatively, for some measurements, unspecifically stuck MyOne superparamagnetic beads were used as reference beads. For liquid exchange in the flow cell, the top coverslip was prepared with two openings with a 1 mm radius. Flow cells are assembled by connecting the two coverslips using a layer of Parafilm (Carl Roth), with a central 50 μL channel cut out, and sealed by heating on a hot plate at 100 °C for approximately 20 s.

After flow cell assembly, 100 μL of 100 μg/mL anti-digoxigenin (Abcam) in 1× PBS are introduced and incubated for at least one hour. The flow cell is passivated using 100 μL of 250 mg/mL bovine serum albumin (Carl Roth) for one hour to counter non-specific interactions. The bead-coupled DNA constructs are introduced in the flow cell and allowed to bind via multiple digoxigenin:anti-digoxigenin interactions to the bottom coverslip during a 30 min incubation.

### Magnetic tweezers setup

A custom-built single-molecule magnetic tweezer setup as described previously was used to perform experiments on DNA (39,40). The setup employs two 5 × 5 × 5 mm^3^ permanent magnets (W-05-N50-G; Supermagnete) with a 1 mm gap placed in a vertical configuration (23). A DC translational motor (M-126.PD2, Physik Instrumente) was used to control the distance between magnets and the flow cell, together with a rotational motor (C-150.PD, PI) to control the rotation of the magnets. We used a LED (69647; Lumitronix LED Technik) for illumination. A 40× oil-immersion objective (UPLFLN 40×; Olympus) and a CMOS sensor camera with 4096 × 3072 pixels (12M Falcon2; Tele-dyne DALSA) were used to image a field of view of 400 × 300 μm^2^. Images recorded at 58 Hz were transferred to a frame grabber (PCIe 1433; National Instruments). The images were tracked in real time using custom tracking software (Labview, National Instruments) to extract the beads’ (*x, y, z*) coordinates (41,42). The objective was positioned on a piezo stage (PifocP726.1CD, Physik Instrumente) to create a look-up table for tracking of the beads’ z-position. The look-up table was generated over a range of 20 μm, with a step size of 100 nm. Labview routines described previously were used for set up control and bead tracking (42). For measurements at different DMSO concentration, the tracked positions using the look-up table (LUT) were corrected for the change in index of refraction with DMSO concentration (Supplementary Figure S1).

### Magnetic tweezers measurements

Prior to measuring, the DNA tethered beads are subjected to force and torque to detect the binding to multiple tethers and to test for torsional constraint. First, negative turns are applied under high tension (*F* = 5 pN). At a high tension, rather than buckling to form plectonemes, melting of DNA occurs upon applying negative turns (43) (corresponding to underwinding of the DNA), preventing a decrease in extension. However, in the case of multiple double-stranded DNA tethers attached to the same bead, a decrease in extension is observed as the tethers braid upon rotation of the magnetic bead. Beads bound to multiple tethers are not analyzed further. To test for breaks or nicks in the DNA tether, positive turns are applied at low tension (*F* = 0.5 pN). In the case of torsionally constrained DNA, the tethers will overwind and form plectonemes, consequently decreasing the extension. In contrast, torsionally unconstrained tethers will not overwind and no plectonemes are formed.

First, rotation-extension measurements are performed on torsionally constrained tethers at several forces (0.25, 0.5, 0.8 and 1 pN) by determining the extension while varying the relative turns of the magnet. The linking number is changed in steps of five turns for 0.25 pN and in steps of 10 turns for 0.5 pN and higher; at each force and number of applied turns, the extension is measured for 6 s and averaged. To determine the initial orientation of the tethers, a rotation curve is measured in 1× PBS. While applying a constant force, the magnets are rotated to positive or negative turns, and the extension is monitored. From this, the relative number of turns to achieve maximum extension and thus fully torsionally relaxed tethers (which we designate as zero turns) is determined. Second, the force-extension response is determined by measuring the tether extension while placing the magnets subsequently at 11 positions corresponding to forces between 0.07 – 5 pN.

The effects of different concentrations of DMSO on the rotation-extension and force-extension behavior of DNA are investigated by subsequently changing DMSO concentration and repeating the rotation and force-extension measurements. At each concentration step, 200 − 300 μL (corresponding to 4-6 flow cell volumes) of DMSO solution at defined concentrations are introduced to the flow cell using a peristaltic pump, in order from low to high concentration, unless otherwise noted. We applied a force of 2 pN while flushing to constrain movement of the bead and to prevent changes in bead rotation during fluid exchange.

### DNA preparation for AFM imaging

Linear dsDNA was obtained by cutting the commercially available DNA vector pBR322 (New England Biolabs) with the FastDigest enzyme EcoRI (Thermo Fisher). Following the manufacturer’s instructions, we scaled up the reaction using 5 µg of plasmid DNA in a total volume of 50 µL. The enzymatic reaction was stopped by heat inactivation and the success of the digest was confirmed in a 1% agarose gel using ROTI-Gel stain (Carl Roth) for DNA visualization and 1 kb Plus DNA Ladder (NEB) as a size reference (Supplementary Figure S2). The linearized DNA was purified using a PCR clean up kit (Qiagen). Differing from the manufacturer’s clean up protocol, the DNA was washed in three steps and eluted in two steps of 25 µL 1× PBS after an incubation time of 1 min at 37 °C.

### AFM imaging

For imaging we use a Nanowizard Ultraspeed 2 high speed AFM (JPK) with triangular FASTSCAN A probes (Bruker) with a driving frequency of ∼1400 kHz and a spring constant of 18 N/m. Overview images for analysis were scanned in tapping mode in air over a field of view of 6 × 6 µm^2^ at 512 × 512 pixels and a line scanning speed of 1 Hz. High-resolution images for additional visualization were recorded with 2048 × 2048 pixels in a field of view 1 × 1 µm^2^. Data for end-to-end distances were obtained from 3-5 independent imaging sets, each containing 3-7 images per measurement condition. Area and length of the longest axis parameters were obtained from single molecule analysis of 3 images per condition.

Before surface deposition and imaging, 18 ng of linearized DNA in 1× PBS buffer were incubated on ice with varying concentrations of DMSO (0, 2, 10, 20, 50 (% v/v)) in a total volume of 20 µL. Two independently prepared batches of linearized DNA were used for imaging and gave consistent results (Supplementary Figure S3). A comparison of incubation times (Supplementary Figure S4) found no significant effect and we chose an incubation time of 20 min for all conditions. After incubation in buffer with varying concentrations of DMSO, the samples were drop casted onto freshly cleaved poly-L-lysine (0.01% (w/v)) coated mica (TED PELLA) and incubated for 30 s, extensively rinsed with ultrapure water, and dried in a gentle stream of pure nitrogen gas (28,44,45).

### AFM image processing and analysis

AFM images were processed with the MountainsSPIP 10 analysis software (Digital Surf) applying line-by-line correction and polynomial background subtraction. End-to-end distances of individual DNA molecules were measured using the manual distance measurement tool of the software by drawing a line between the identified free ends of single DNA molecules (Figure 5A). The end-to-end distance is given by the horizontal distance between the first and the last point of the segment, along the defined path, measured on a horizontal plane. For end-to-end distance analysis we analyzed > 125 individual DNA molecules for each measurement condition. Mean-squared end-to-end distances <*R*^2^> were directly computed from the data. In addition, we fit the distributions of end-to-end distances *r* obtained by AFM imaging using the probability function of a polymer chain in three dimensions (46,47):

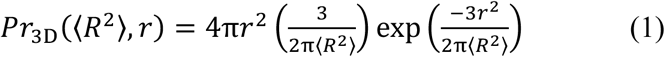

We note that the 3D Gaussian chain model is only an approximation to the situation in the AFM imaging. Therefore, Equation 1 is primarily used as a convenient expression that fits the data (see also the next section).

In addition to the end-to-end distances, we obtained shape parameters for the area and the long-axis length of the DNA, again from the MountainsSPIP 10 software, using the manual shape analysis tool to draw contours around single DNA molecules manually. The area parameter is given by the projected horizontal area enclosed by the outline of the molecule. The long-axis length parameter is calculated by the software from the molecule’s oriented bounding rectangle, which is parallel to the longest axis of the particle. With the area and long-axis length parameters, we can evaluate the compaction of every single DNA molecule that we could assign in the images and are not restricted to molecules with visible free ends, as it is the case in the end-to-end distance analysis. The shape parameter analysis was applied to every single DNA molecule identified in a subset of 3 images per condition.

### Models and scaling of the end-to-end distances of DNA

We use results from the statistical mechanics of semi-flexible polymers to connect the DNA parameters determined from the MT force-extension measurements to the AFM results. The mean squared end-to-end distance in 3D has been shown to be related to the bending persistence length *L*_P_ and contour lengths *L*_C_ via (48):

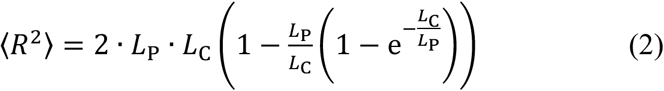

Equation 2 describes the mean squared end-to-end distance in 3D, while AFM images are recorded at a surface. To take into account the fact that AFM images are recorded in 2D, two limiting cases are often considered (49): (i) full 2D equilibration, where the chain completely equilibrates at the surface, (ii) and kinetic trapping, where the chain immediately “sticks” to the surface. Full 2D equilibration at the surface is expected to result in an equilibrated 2D chain for which *L*_P_ in Equation 2 is replaced by 2×*L*_P_. Therefore, a 2D equilibrated chain is predicted to have an approximately 2-fold larger value for <*R*^2^> than in 3D. Conversely, the kinetic trapping regime has been approximated by a simple, mathematical projection of the 3D chain onto 2D (48), in which case Equation 2 is multiplied by 2/3. Importantly, both limits are approximations and real imaging conditions are likely between these extremes. It has been shown that our imaging conditions (deposition on PLL-mica) do not lead to 2D equilibration (49,50), but that the DNA conformations are, at least approximately, representative of the DNA structure in solution. Further, all three scenarios for <*R*^2^> (Equation 2 for 3D, full 2D equilibration, and the 3D-to-2D-projection) predict the same scaling of the mean squared end-to-end distance <*R*^2^> ∼ *L*_P_ · *L*_C_.

### Coarse-grained Monte Carlo simulations of DNA

We performed Monte Carlo simulations of a modified worm-like chain (WLC) model, employing a simulation procedure similar to what was used previously (32,51–53). The DNA is modeled as a chain of rigid bodies represented by right-handed reference frames (triads) akin to the rigid base pair model (53–56). The molecule is assumed to be inextensible, i.e. we consider a constant discretization length of *a =* 0.34 nm, corresponding to the length of one base pair (57,58). Individual configurations are parametrized in terms of rotation vectors that transform adjacent triads into one another (for a detailed discussion see (53)). The three components of the vector, expressed in the local material frame, are identified with two local bending degrees of freedom (Ω_1_ and Ω_2_) and twist (Ω_3_). We note that our simulations have freely rotating ends, unless otherwise noted, such that the twist does not influence the results.

Although this description allows for the consideration of sequence-dependent structure and elasticity (53–55), we ignore sequence effects and consider two types of base pair steps: duplex and bubble. Duplex steps are parametrized with a bending persistence length of *L*_P_,_duplex_ = 43 nm and a twist persistence length *C*_duplex_ = 100 nm. For bubble steps, which represent local defects or perturbations of the DNA helix induced by DMSO, we consider various bending stiffnesses *L*_P_,_bubble_ with values ranging from 0.5 to 5 nm, which span the range of values reported for the persistence length of single-stranded DNA (59,60) and DNA loops (61). Similarly, bubble steps are modeled with a reduced twist stiffness of *C*_bubble_ = 25 nm. Additionally, for duplex steps, we impose an intrinsic twist of *ω*_0,duplex_ = 34°, reflecting the natural helical winding of the molecule. For bubble steps, we introduce two modified values of intrinsic twist, *ω*_0,bubble_ =17° and 0°, corresponding to half- and full-unwinding of the affected segment, respectively. These elastic constants are incorporated into the elastic Hamiltonian of a N+1 base pair molecule as:

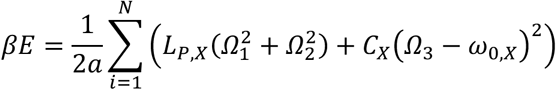

where *β* is the inverse temperature, i.e. *β* =1/*k*_B_T where *k*_B_ is Boltzmann’s constant and *T* the absolute temperature, and X is either equal to bubble or duplex, depending on the type of the segment.

Simulations were conducted under varying conditions to explore the potential distribution and clustering of bubble segments. Bubble domains were modeled to vary in size, ranging from individual segments to full phase separation, where all bubble segments are clustered into a single domain. Intermediate bubble sizes of 1, 3, 10, 25, 100, 250 segments and full phase separation were simulated to capture different degrees of bubble organization. For each bubble size, simulations are initiated with a randomly chosen but fixed distribution of bubbles. To account for variations in bubble positioning, we performed 20 independent simulations with different randomly chosen bubble locations for each set of simulation parameters. Since we do not know *a priori* how many bubbles there are and how their number depends on DMSO concentration (and possibly force and torque etc.), we treat the fraction of bubbles *f*_bubble_ as an adjustable parameter. We simulate chains over a wide range of bubble fractions *f*_bubble_ = 0 – 0.2, to identify which parameters are consistent with experimental observations. Ensembles of configurations are generated with a combination of cluster moves (crankshaft and pivot moves (32,52)). For the tweezer setup simulation, an additional force-dependent term βf⃗ ⋅ ^?^R⃗, where *R* is the end-to-end distance of the molecule, is introduced to the elastic energy (51). Each simulation was run for 10^8^ Monte Carlo steps, which is much longer than the typical convergence sampling of 10^6^ – 10^7^ steps.

## RESULTS AND DISCUSSION

To determine the effects of DMSO on the mechanical properties of DNA, in particular to test whether the presence of DMSO induces lengthening, unwinding, or a change in the bending stiffness of DNA, we performed single-molecule MT measurements (Figure 1; Materials and Methods). In a first set of experiments, we measured the DNA extension at different levels of applied force for torsionally relaxed DNA. In a second set of measurements, we probed the response of DNA to systematic over- and underwinding at constant forces. We then probe DNA conformation in the presence of different DMSO concentrations in the absence of external forces by AFM imaging. Finally, we present coarse-grained Monte Carlo simulations to rationalize our findings from a microscopic view.

**Figure 1.**
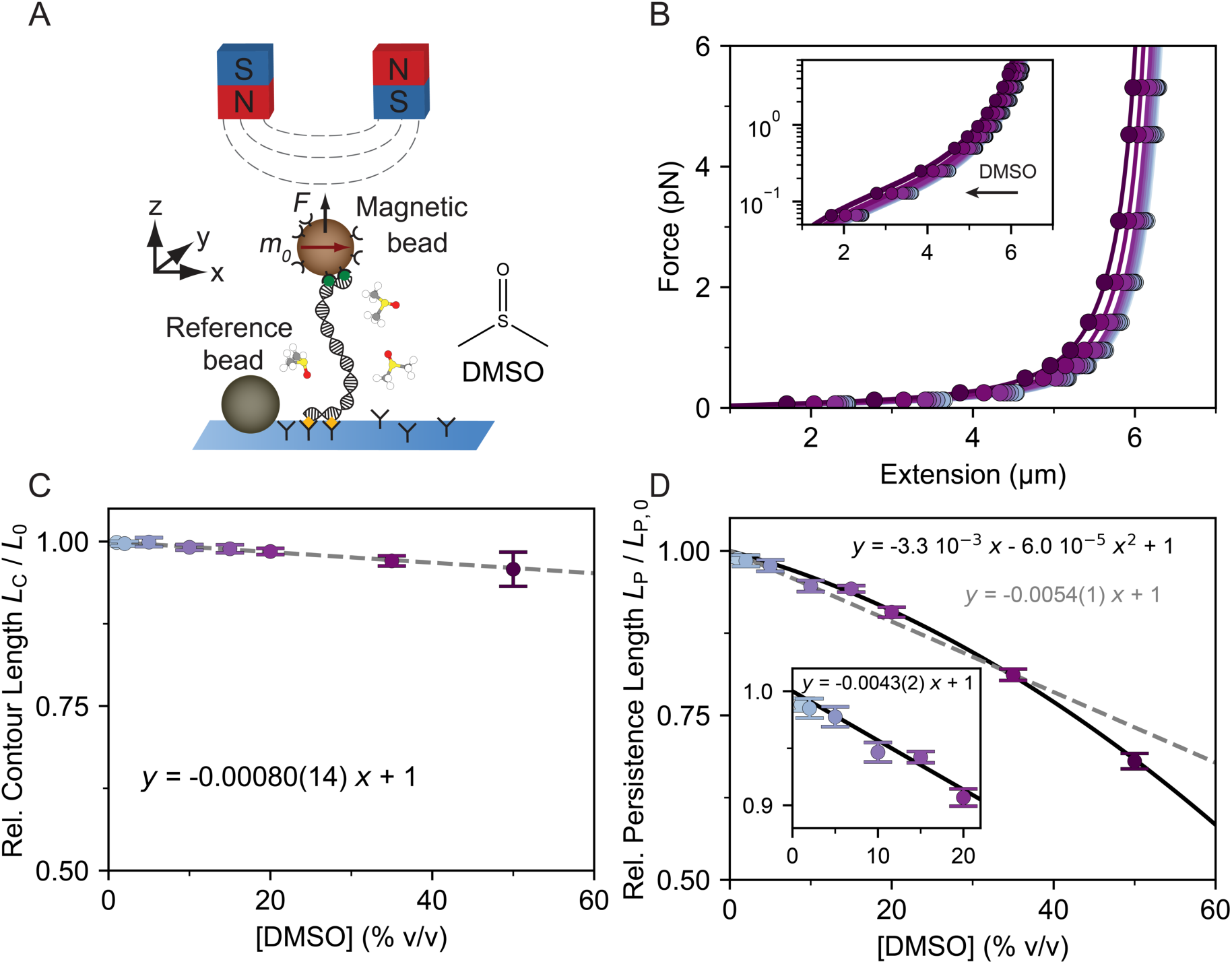
Magnetic tweezers force-extension measurements reveal a linear decrease in contour and bending persistence length with increasing DMSO concentration. A) Schematic of the magnetic tweezers measurement. DNA molecules are tethered between a surface and magnetic beads in a flow cell. Permanent magnets above the flow cell enable the application of stretching forces *F* and torques. B) Force-extension measurements in the presence of increasing concentration of DMSO for 20.6 kbp DNA in PBS buffer. The DMSO concentrations are (from light blue to purple) 0, 1, 2, 5, 10, 15, 20, 35, 50% (v/v). Solid lines are fits of the worm-like chain (WLC) model, treating the contour length *L*_C_ and bending persistence length *L*_P_ as fitting parameters. C) Contour length *L*_C_ as a function of DMSO concentration was obtained from WLC fits as shown in panel B. Data are normalized to the fitted contour length in the absence of DMSO (i.e. in PBS buffer), which is 6.9 ± 0.2 µm for our 20.6 kbp DNA construct. D) Bending persistence length *L*_P_ as a function of DMSO concentration was obtained from WLC fits as shown in panel B. Data are normalized to the fitted persistence length in the absence of DMSO (i.e. in PBS buffer), which is 43 ± 6 nm. The black line is a second-degree polynomial fit to the data. Inset shows a linear fit to the data up to 20% DMSO with a (0.43 ± 0.02)% decrease in *L*_P_ per vol%. Data in panels C and D are the mean ± SD and mean ± SEM from at least 8 independent molecules, respectively. Fitted parameters are indicated as insets.

### DMSO causes a gradual decrease of DNA bending persistence length

To investigate the effect of DMSO on DNA length and bending stiffness, we used MT to perform stretching experiments for increasing DMSO concentrations (0-60%) (Figure 1A). We first performed force-extension measurements on torsionally unconstrained DNA at stretching forces ≤ 5 pN: in this range the response of DNA is dominated by entropic stretching elasticity (62–64). The inextensible WLC model (62–64) describes the resulting force-extension curves well (Figure 1B) and for DNA in PBS without DMSO we find a contour length of *L*_C_ = 6.9 µm ± 0.2 µm and a persistence length of *L*_P_ = 43 nm ± 6 nm (mean ± standard deviation from 86 molecules) in excellent agreement with previous measurements (23,64,65).

Next, we measured force-extension curves after addition of increasing amounts of DMSO (Figure 1B). From fits of the WLC model, we find a systematic decrease in the contour length (Figure 1C) and bending persistence length (Figure 1D) with increasing DMSO concentrations. The observed decrease in contour length is approximately linear and very small, < 5% even at 50% DMSO. A linear fit gives a decrease in the apparent contour length of 0.08% per vol% DMSO, but we note that the change is essentially within experimental error.

In contrast, we find an overall still moderate, but systematic decrease in DNA bending stiffness for increasing DMSO concentrations (Figure 1D). For low and intermediate concentrations of DMSO, the decrease in bending persistence length with increasing DMSO concentration is linear and from a linear fit up to 20% DMSO (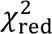 = 1.3) we find a (0.43 ± 0.02)% (the error is obtained from the covariance matrix of the fit) decrease in bending persistence length per vol% DMSO (Figure 1D, inset). For higher DMSO concentrations a larger dependence is observed (Figure 1D). The entire concentration range is less well described by a linear dependence, a linear fit up to 50% DMSO gives a (0.54 ± 0.01)% decrease per vol%-DMSO with a 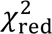 = 7. We find that a second-degree polynomial provides a good description of the data (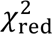 = 1.3; Figure 1D).

Control measurements show that the change in bending persistence lengths is not due to the (small) reduction in ionic strength upon addition of DMSO, since results within experimental error are obtained if the ionic strength and buffer concentration are kept constant upon addition of DMSO (Supplementary Figure S5). In addition, a decrease in ionic strength (upon dilution) would be expected to result in an increase and not decrease in bending persistence lengths (66–69).

### DNA stability and collapse in high DMSO concentrations

We tested the stability of DNA in moderate concentrations of DMSO by monitoring the force response of DNA for a long period of time (1.5 h) in 20% DMSO (Figure 2A). The extension time traces show extension changes in response to changes in applied force, as expected from the WLC behavior discussed above, but exhibit no time dependence otherwise. In particular, no sudden decrease in extension or compaction of the DNA is observed in the range of forces investigated (0.07 – 5 pN). The extension at a given force remains constant and is reproducibly achieved in repeat measurements lasting > 1 h (Figure 2B).

**Figure 2.**
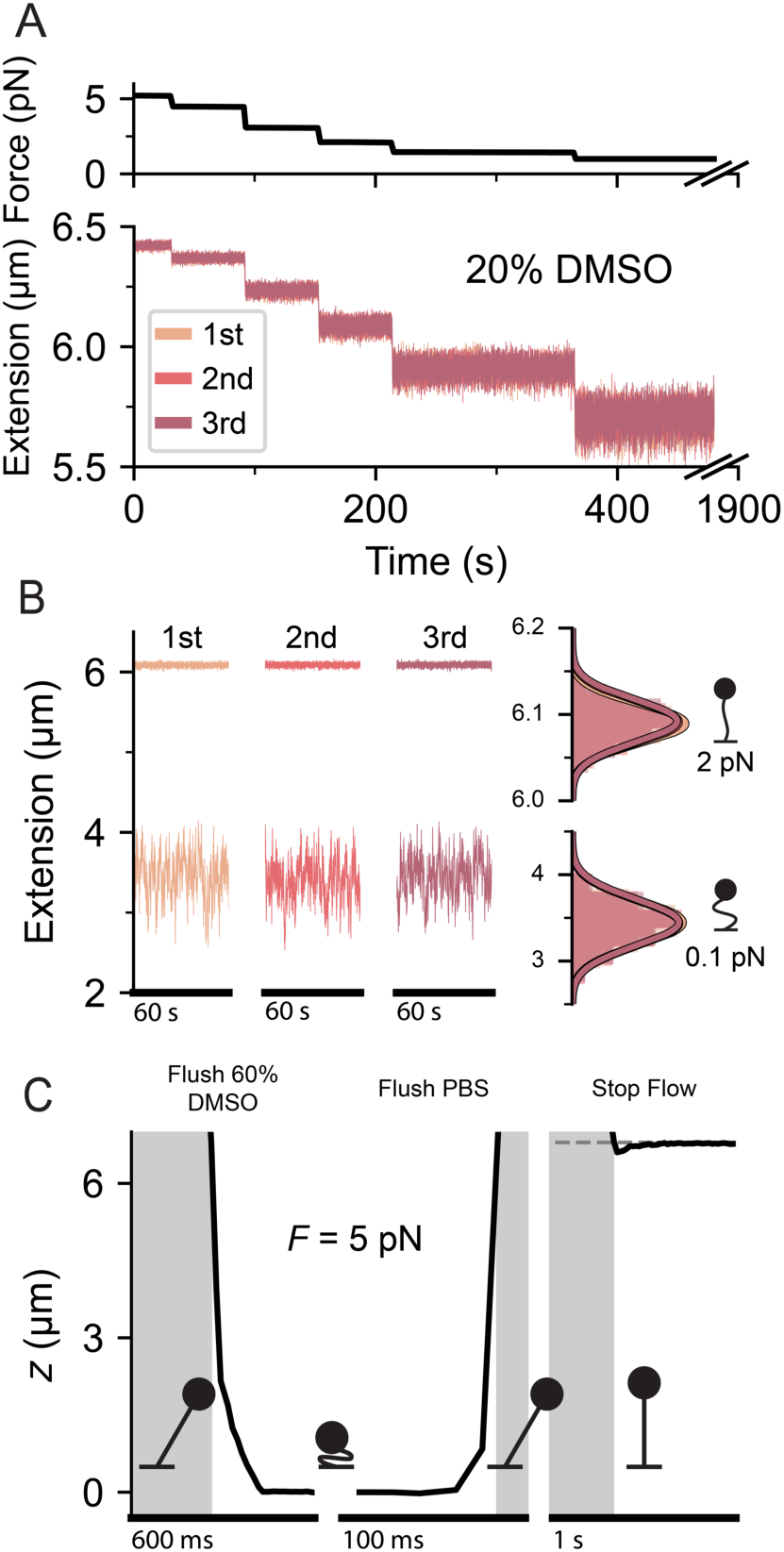
Magnetic tweezers measurements reveal the effect of DMSO on the stability of DNA. A) Force response of our 20.6 kbp DNA construct in 20% (v/v) DMSO. Top: Forces applied to the molecule vs. time. Bottom: Extension of the molecule vs. time. The data shown are from 3 subsequent force scans (indicated by the different colors), recorded over a period of time of 1.5 hours. The response of the DNA remains highly consistent throughout the subsequent force cycles, such that traces are superimposable. B) 60 s segments from extension time traces at the forces indicated on the right (0.1 or 2 pN) in 20% DMSO. The full extension time traces at 0.1 pN are significantly longer (300 s) than the segments shown. The behavior of the molecule remains unchanged, even after being at low force (0.1 pN) for a long period of time (5 min). Right inset shows the distribution of measured extensions for the total time trace at the given force. C) Extension-time traces of DNA in the presence of 60% DMSO. DNA collapses against a constant 5 pN stretching force as a 60% DMSO solution is flushed into the MT flow cell. The DNA collapse is reversed upon flushing PBS buffer, when the molecule extends again to its initial length. The horizontal grey dashed line is the measured extension at 5 pN stretching force in PBS prior to DNA collapse. The scale bars indicate the duration of the segments. The vertical grey boxes indicate times when tracking of the magnetic bead is disrupted due to flushing for buffer exchange. Total flushing time of both 60% DMSO and PBS was ∼8 min.

To investigate the behavior of DNA in the presence of very high concentrations of DMSO, we increased the concentration to 60% DMSO. Under these conditions, the DNA collapses against stretching force of 5 pN into a compact, condensed state (Figure 2C). This collapse is reversible: upon reintroduction of the initial buffer condition, the DNA returns to the extension measured prior to collapse in absence of DMSO (Figure 2C, right panel, grey dashed line). This collapse is likely related to a decrease in solvent quality of mixtures with high fractions of DMSO, as has been observed for lower DMSO concentrations in the presence of high concentrations of divalent metal ions (70) and in organic solvents and alcohols (71). Since the emphasis in this work is on the properties of DNA under low and intermediate concentrations of DMSO, as are e.g. typically used as additives in enzymatic assays, we limited our measurements to ≤ 50% DMSO in the remainder of this work.

### DMSO has only minor effects on DNA twist

Effects on the twist *Tw* of DNA can be monitored by control of the molecule’s linking number *Lk*. In the case of torsionally constrained DNA, the linking number *Lk* can be directly controlled in the MT by rotation of the external magnets. We find the magnet orientation at which DNA is torsionally relaxed in PBS buffer in the absence of DMSO, which corresponds to the intrinsic helicity *Lk*_0_ of dsDNA and which we refer to as zero turns, by systematically over- and underwinding the molecule (Figure 3A). At low applied stretching forces (*F =* 0.25 pN), an approximately symmetric response upon under- and overwinding is observed, with buckling transitions for negative and positive turns. Past the buckling transitions, a linear decrease in extension is observed upon further rotation of the magnets, corresponding to the formation of positive or negative plectonemic supercoils of the DNA strands (21,24). We quantify the change in the twist of DNA by extrapolating the slopes in the plectonemic regime (72), which provides the intersection points, which indicate the points where the molecule is torsionally relaxed. In contrast to the contour and persistence length, the twist of DNA is affected much less by increasing DMSO concentrations (Figure 3B). The intrinsic helicity in PBS remains unchanged, within experimental error, below 20% DMSO, which suggests that there are no large open regions or extended melting bubbles. Above this concentration we observe a small decrease in the DNA twist, by (0.22 ± 0.08) degrees per base pair for 50% DMSO. We note that a reduction of DNA twist at high DMSO concentrations is both qualitatively consistent with changes of the double-helix due to reduced electrostatic screening and also with local melting of the DNA. DNA twist decreases with decreasing salt concentration (73,74); DMSO has a lower dielectric constant than water (≈47 for pure DMSO vs. ≈80 for water), which implies that electrostatic interactions would be increased with increasing DMSO concentrations. At the same time, local DNA melting is associated with unwinding of the helix (24,75,76). Both effects are predicted to cause DNA unwinding.

**Figure 3.**
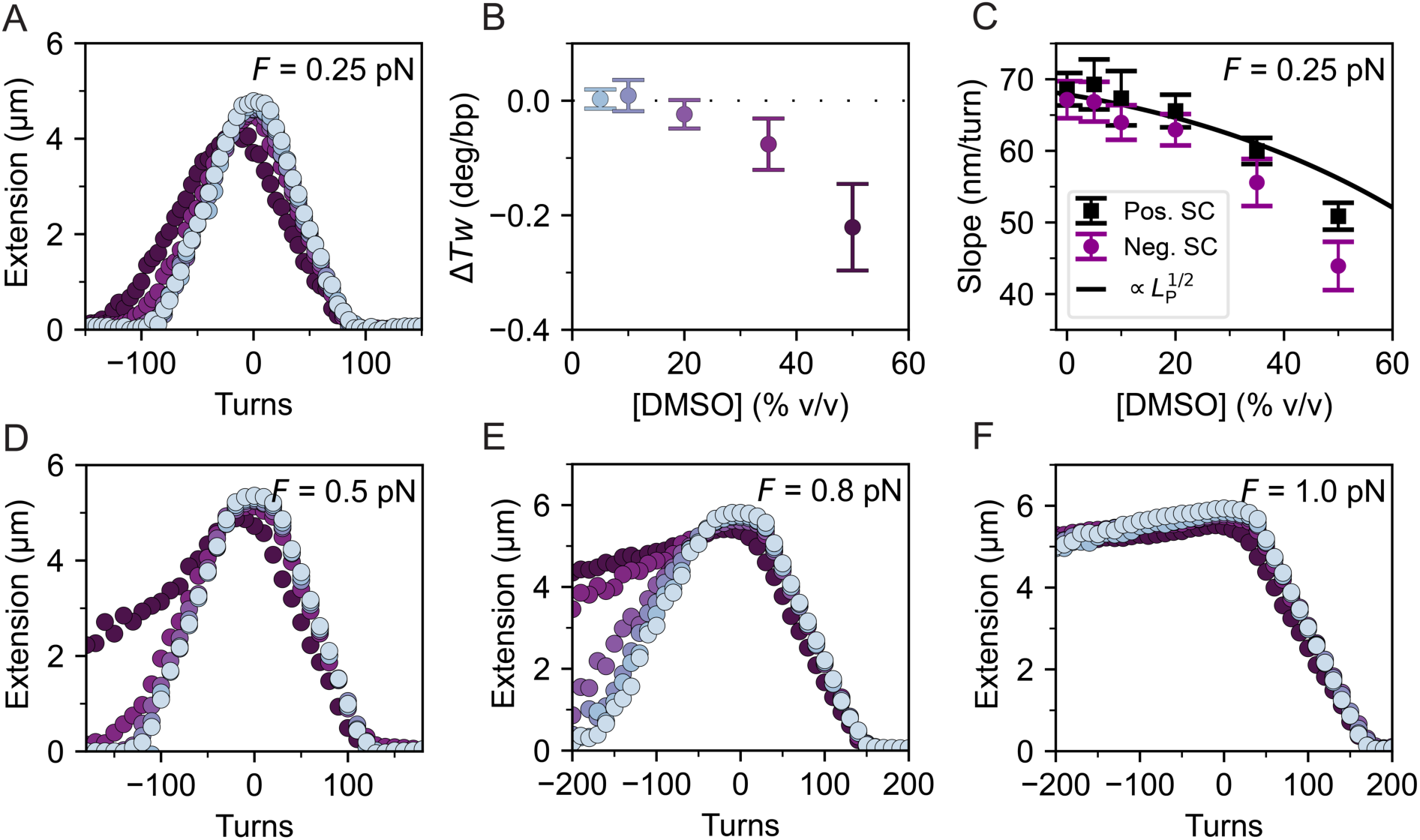
Magnetic tweezers rotation extension measurements show that increasing DMSO concentrations facilitates melting but have only a small effect on DNA twist. A) Rotation-extension measurements of a 20.6 kbp DNA in the presence of increasing concentration of DMSO (from light blue to purple; same color code as Figure 1). B) Change in DNA twist as a function of DMSO concentration determined from the centers of rotation curves as in panel A. The twist is measured relative to DNA in the absence of DMSO (i.e. in PBS buffer). C) Slopes of the rotation-extension curves in the plectonemic regime at *F* = 0.25 pN for positive and negative turns (relative to relaxed DNA). Data in panels B and C are the mean ± standard deviation from at least 5 independent molecules (at least 11 for DMSO concentrations ≤ 10%). The solid line is the square root of the quadratic fit to the *L*_P_ / *L*_P,0_ data from Figure 1E, normalized to the slope in the absence of DMSO (i.e. in PBS buffer). A dependence of the slope in the plectonemic regime on *L*_P_^1/2^ is predicted by models of DNA supercoiling (22,32,77,78). D-F) Rotation-extension curves for 20.6 kbp DNA as a function of DMSO concentration at forces of 0.5 pN (D), 0.8 pN (E), and 1.0 pN (F). While curves at low forces (e.g. panel A) are approximately symmetric around zero turns, the rotation curves at higher forces and higher DMSO concentration are asymmetric, exhibiting lower (absolute) slopes for negative turns than at positive turns, due to torque-induced melting. It is apparent that the onset of torque-induced melting occurs at lower forces for higher DMSO concentrations.

From the rotation curves at low forces, *F =* 0.25 pN, we quantify the slopes of turns vs. extension past the buckling points, i.e. in the negative and positive plectonemic regimes (Figure 3C). We find that at this low force, the slopes in the negative and positive plectonemic regime remain equal, within experimental error, for all DMSO concentrations except for 50% DMSO. In addition, the dependence on DMSO concentration can be accounted for by the change in bending persistence length (Figure 1D) using the scaling proportional to *L*_P_^1/2^ that is predicted by models of DNA supercoiling (22,32,77,78) (Figure 3C, black line).

### DMSO facilitates torque-induced melting

Next, we measured the effect of DMSO on rotation-extension curves at different forces (0.25, 0.5, 0.8, 1 pN, Figure 3), to monitor the increase in DNA melting by DMSO using MT. In the absence of DMSO, the rotation-extension curves of bare DNA are symmetric at forces below ≈1 pN, presenting buckling points for both negative and positive turns. At forces ≥ 1 pN, DNA buckles when overwound, but underwinding results in only a small decrease in extension, due to torque induced melting, such that further unwinding is accommodated by DNA melting (24,43). At a force of ≈ 0.8 pN in PBS buffer, the DNA rotation curves are still essentially symmetric, but one notices increased fluctuations in the extension (Figure 3E and (32)).

In the presence of increasing concentrations of DMSO, the DNA rotation curves start to be asymmetric at lower forces compared to PBS buffer only, suggesting an increase in torque-induced melting in the presence of DMSO. This is consistent with the view that DMSO weakens the hydrogen bonds between the bases, which would correlate to destabilization of the double-stranded DNA form observed in the MT, in line with the reported decrease in melting temperature of DNA in the presence of DMSO (16,79).

### AFM imaging reveals a gradual compaction of DNA upon addition of DMSO

MT measurements constrain the ends of the DNA and stretch the molecule, which might potentially suppress the effects of compaction, local melting, or kinks on its conformations. Therefore, we performed AFM imaging experiments to visualize the conformations of linear dsDNA in the absence of external forces and torques after surface deposition (49,80,81) in near-physiological PBS buffer with varying DMSO concentrations ranging from 0% to 50% (Material and Methods). Our AFM images, recorded in tapping mode in air, clearly visualize individual DNA molecules, showing the typical “worm-like chain” conformations of semi-flexible polymers and suggest a small, but systematic compaction of the DNA conformations with increasing DMSO concentration (Figure 4).

**Figure 4.**
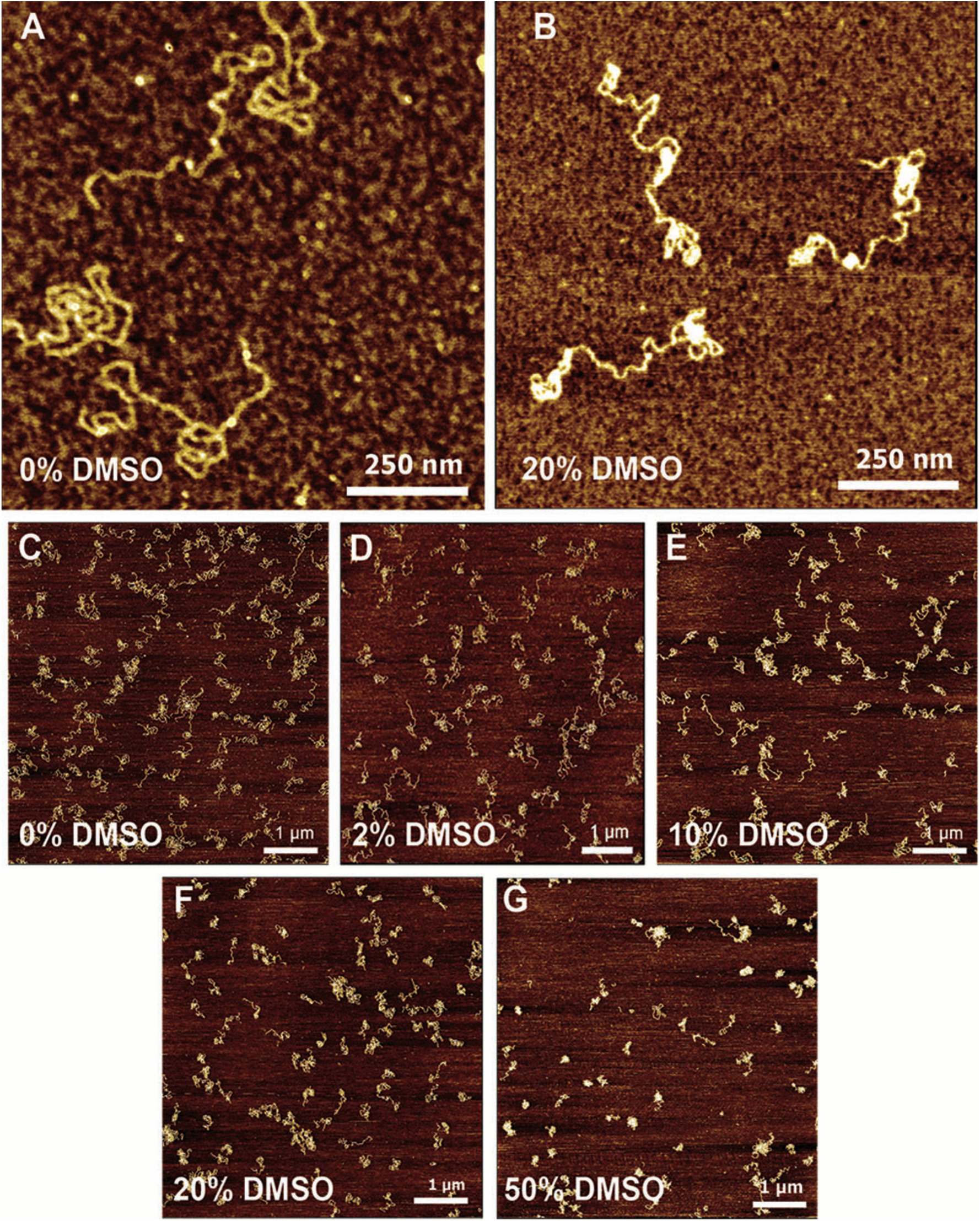
AFM imaging probes DNA in the presence of DMSO. Topographic images of linearized pBR322 DNA on PLL-mica measured in air without DMSO (A and C) and with DMSO concentrations ranging from 2% to 50% (B and D-G). It is visible that DNA moderately compacts in high DMSO conditions (≥ 20%) while differences in DNA structure and compaction are not distinguishable by eye for DMSO concentrations below 20%. High resolution images of DNA are shown for PBS only (A) and for 20% DMSO (B).

To quantify the shape and compaction of individual DNA molecules, we quantified three parameters using the MountainsSPIP 10 analysis software (Materials and Methods): (i) We obtained end-to-end distances using the manual distance analysis tool by drawing straight lines between the free ends of single DNA molecules (Figure 5A). The end-to-end distance is a commonly used parameter to analyze the size of linear polymers in solution. End-to-end distances give an indication of the structural behavior of DNA and with clear predictions from polymer physics and can be linked to parameters from magnetic tweezer experiments via <*R*^2^> ∼ *L*_P_ · *L*_C_ (Equation 2, Material and Methods). (ii, iii) To evaluate and confirm the compaction of DNA without the necessity to identify the ends, which can only be analyzed for ∼40-50% of the molecules (Supplementary Figure S6), we additionally obtained area and long-axis parameters from the AFM images. Using the manual shape analysis tool of the MountainsSPIP10 software, we drew contours around individual DNA molecules (Figure 5A) and obtain the output parameters from the software. While less readily linked to polymer theory, the area and long-axis length parameters provide a control to quantitate systematic changes of DNA conformations upon increasing DMSO concentrations.

**Figure 5.**
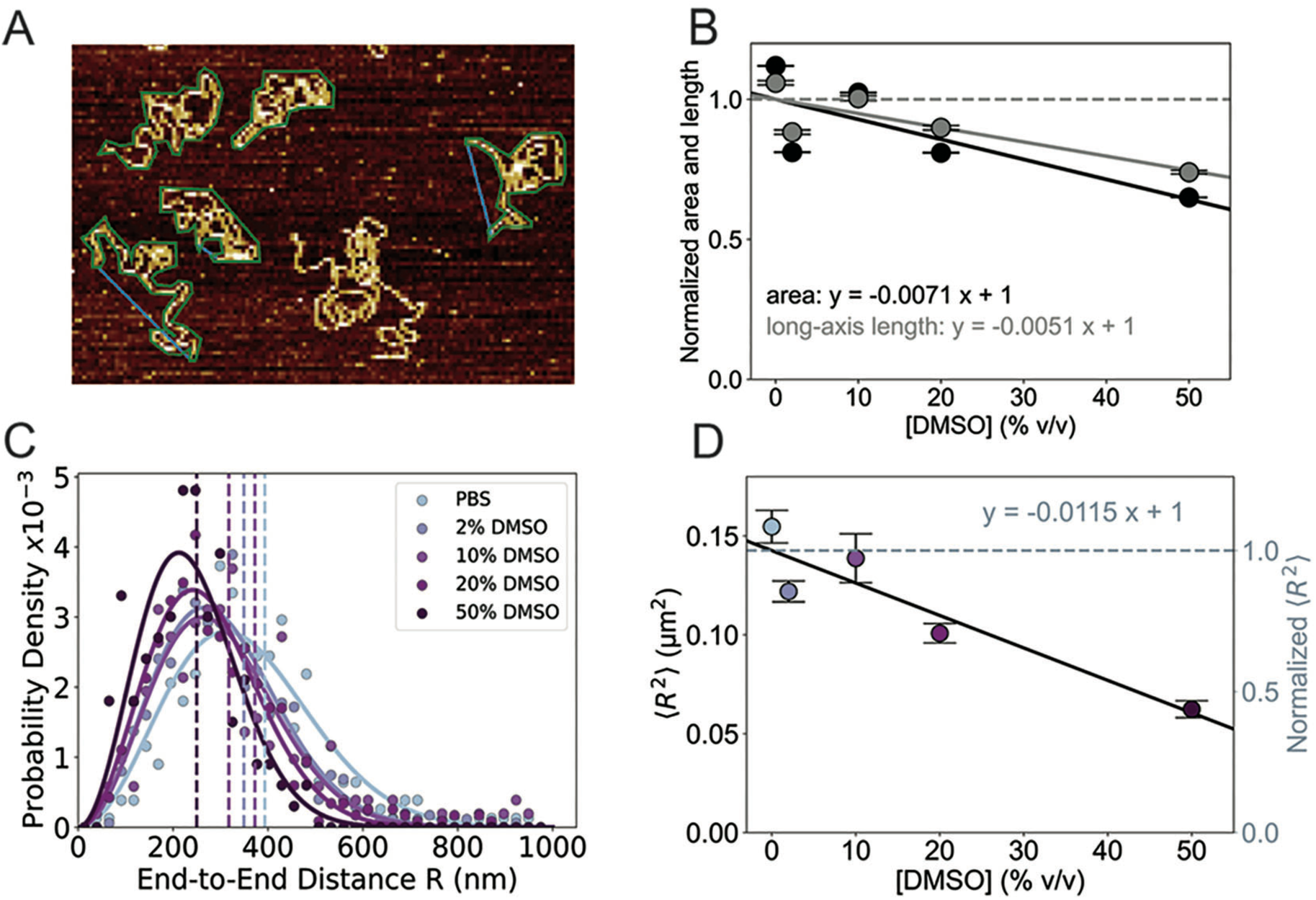
Quantification of AFM images reveals a moderate compaction of DNA in the presence of DMSO. A) Topographic AMF image of linearized pBR322 DNA (4361 bp) in PBS condition on PLL-mica measured in air. End-to-end distances were measured manually using the manual measurement distance tool of the MountainsSPIP 10 software and are indicated by blue lines in the image. Green contours around DNA molecules show how we used the manual shape analysis tool of the MountainsSPIP 10 software to obtain shape parameters like area and long-axis length. B) Long-axis length (shown in grey) and area (shown in black) parameters as a function of increasing DMSO. The data are normalized to the 0% DMSO value from the fit. The solid lines indicate a linear fit with a negative slope corresponding to a decrease in area of 0.71% (black line) and long-axis length of 0.51% (grey line) per %-DMSO. C) Histograms of DNA end-to-end distances with increasing concentrations of DMSO (from 0%, blue to 50%, purple) and fits of the end-to-end distribution of a Gaussian chain (solid lines in the corresponding colors). RMS values are indicated as colored dashed lines and show a trend of decreasing end-to-end distance of the DNA upon increasing DMSO concentration. D) Mean-squared end-to-end distances *<R^2^>* as a function of increasing DMSO concentration. The right axis shows values relative to the 0% DMSO value from the fit. The black line indicates the linear fit with a negative slope corresponding to ∼1.2% decrease of mean-squared end-to-end distance per %-DMSO. Symbols and error bars in panels B and D are the mean ± SEM.

All three parameters show similar trends, namely a moderate but gradual decrease in size upon increasing DMSO concentrations (Figure 5). The area of the molecules shrinks to 58% of its area in the 50% DMSO condition in comparison with no DMSO treatment and decreases approximately linear with 0.71% per %-DMSO over the whole range. The mean long-axis length decreases to 70% of its length at 50% DMSO with 0.51% per %-DMSO, indicating moderate systematic compaction of the DNA molecules (Figure 5B). The effect of DMSO on the end-to-end distances <*R*^2^> is described in detail in the next section.

### DMSO induces a gradual decrease of the end-to-end distances

To quantify the DNA conformations visualized by AFM imaging, we identify the ends of the DNA molecules and measure the corresponding end-to-end distances (Material and Methods; Figure 5A). While the end-to-end distances could not be traced for all molecules visible in the AFM images (∼ 40-50%, Supplementary Figure S6), in particular due to obscured or overlapping ends, we obtained a similar number of traced molecules for each DMSO condition (≥ 125 molecules per condition). The resulting end-to-end distances distributions show the typical positively-skewed distributions previously observed (49) for DNA by AFM imaging and are well-described by the Gaussian chain model shown in Equation 1 (Figure 5C and Supplementary Figure S7). The distributions shift to smaller end-to-end distances with increasing DMSO concentrations and we find that the distributions for all investigated DMSO concentrations are significantly different from the PBS only conditions, as assessed both using two-sample Kolmogorov-Smirnov tests to compare the full distributions and using two-sample two-tailed *t*-tests to compare the means (Supplementary Table S1). Overall, we observe a moderate decrease in mean-squared end-to-end distance <*R*^2^> with increasing DMSO concentrations (Figure 5D), which is reasonably well described by a linear fit. Fitting a straight line, we obtain a slope corresponding to a change by 1.2% decrease in <*R*^2^> per %-DMSO. Using the <*R*^2^> values obtained from fits of the Gaussian chain model (Equation 1) instead of the directly computed <*R*^2^> values gives similar results (Supplementary Figure S8).

### Comparison of MT and AFM results for DNA conformations with increasing DMSO

We can use results from the theory of semi-flexible polymers to connect the results of AFM and MT force spectroscopy (Materials and Methods). A complication in the interpretation of AFM images is that they are recorded for molecules deposited on a surface and that surface deposition conditions can influence what conformations are observed. It has been shown that our imaging conditions (deposition on PLL-mica) do not lead to 2D equilibration (49,50). For simplicity, we use the 3D prediction (Materials and Methods, Equation 2) for the DNA scaling and find reasonable agreement with the experimentally determined values: In PBS only, we measure experimentally <*R*^2^>^1/2^ = 393 nm ± 9 nm (mean ± standard error of the mean), while the 3D model predicts <*R*^2^>^1/2^ = 352 nm ± 24 nm, where we assume a contour length of 0.34 nm/bp and use *L*_P_ and its propagated error from the MT experiments.

Combining the dependences of *L*_P_ and *L*_C_ on DMSO from the MT data, we predict a reduction in <*R*^2^> by 0.62% per %-DMSO, which is smaller but of similar magnitude compared to the experimentally determined slope of 1.2% decrease of <*R*^2^> per %-DMSO (Figure 5D) and the change in <*R*^2^> values determined from the Gaussian fit (Equation 1) that decreases by 0.99% per %-DMSO (Supplementary Figure S8). Similarly, using the *L*_P_ and *L*_C_ determined from MT measurements directly to predict <*R*^2^> at different DMSO concentrations via Equation 2 finds values close to the data determined from AFM imaging (Supplementary Figure S9). In summary, we find an overall consistent view of the changes in DNA mechanics upon addition of DMSO from MT force spectroscopy and AFM imaging analysis, which probes DNA in the absence of applied forces or torsional constraints. The moderately larger effects seen by AFM imaging compared to MT force spectroscopy might indicate that applied forces reduce the impact of DMSO on DNA mechanics.

### Coarse-grained Monte Carlo of DNA with flexible segments

To investigate the observed effects of DMSO from a mechanistic and more microscopic perspective, we turned to coarse-grained Monte Carlo simulations and introduce a minimal model that features local defects, which we refer to as “bubbles”. We model DNA as a discrete worm-like chain (WLC) with two types of inextensible segments each of size 0.34 nm, corresponding to the length of one base pair (Materials and Methods; Figure 6A). One type of segment represents double-stranded DNA with a persistence length of 43 nm (matching the force-extension measurements of DNA in PBS buffer) and the second type of segment represents DMSO-induced defects, which are WLC segments of strongly enhanced flexibility. For simplicity we omit enthalpic stretching and, thus, assume the discretization length to be constant. The bubbles represent regions of local base pair disruption and have significantly lower bending stiffness *L*_P,bubble_ (with *L*_P,bubble_ << *L*_P_), but no intrinsic curvature (which has been considered previously in models of kinked DNA (27)). While the fraction *f*_bubble_ and the location of the bubbles remains fixed throughout each individual simulation, the effect of positional bubble fluctuations is mimicked by repeating simulations for each set of parameters for many different bubble distributions. Furthermore, our and previous (15) experimental data suggest that even in 50% DMSO DNA remains predominantly double stranded, suggesting that *f*_bubble_ is small, i.e. *f*_bubble_ ≪ 1. Likewise, rather than assuming a certain bending stiffness for the bubble segments, we repeat simulations for values of *L*_P,bubble_ ranging from 0.5 nm to 5 nm.

**Figure 6.**
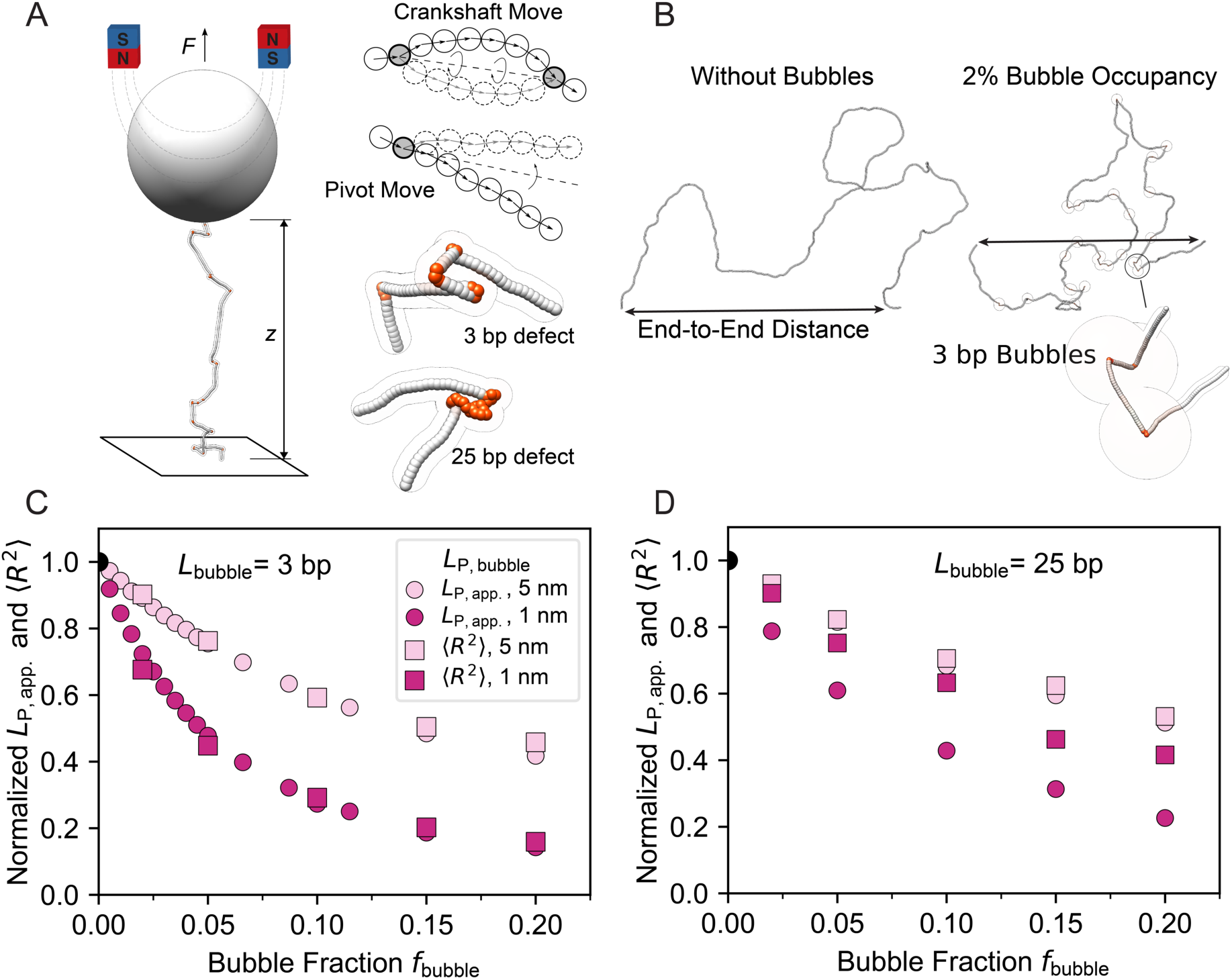
Coarse grained simulations of kinkable DNA under external forces and in free solution. A) Schematic of the coarse-grained MC simulations. The DNA is represented as a chain of beads connected by springs that set the bending stiffness of the chain. In force extension simulations the ends of the chain are constrained and a constant force in *z*-direction is applied, mimicking magnetic tweezers measurements. Crankshaft and pivot moves ensure efficient sampling in the MC protocol (see Methods). Examples of flexible segments or bubbles are highlighted in red. B) Snapshots of coarse-grained MC simulations of 4 kbp DNA in free solution. End-to-end distances are annotated in the plots. C,D) Normalized bending persistence length obtained from fits of the WLC model to simulation data of 8 kbp DNA and normalized mean-square end-to-end distances <*R*^2^> from simulations of 4 kbp DNA in free solution. The apparent bending stiffness decreases systematically with increasing fraction of bubbles and with decreasing persistence length of the flexible segments. <*R*^2^> decreases systematically with increasing fraction of bubbles and with decreasing persistence length of the flexible segments. For bubbles of 3 bp the DNA remains well described by the WLC and obeys the dependence of <*R*^2^> ∼ *L*_P_ · *L*_C_. For larger bubbles, i.e. 25 bp, the DNA is less well described by the WLC for a small persistence length of the flexible segments and shows a decreased dependence of bubble fraction. Similar simulations for other parameters are shown in Supplementary Figures S10-S14.

### Coarse-grained Monte Carlo simulations quantify the effect of DNA bubbles on force-extension measurements and on DNA in free solution

To enable comparison with our MT measurements and AFM imaging data, we simulated the force-extension response of DNA and its conformation in free solution, using the coarse-grained model introduced in the previous sections (Figure 6A,B and Supplementary Figure S10). The simulated force-extension response of fully double-stranded DNA is in excellent agreement with the WLC model and recovers the input value of *L*_P_ from a fit to the simulated data for bare DNA (Supplementary Figure S10 and S11).

We then introduce increasing numbers of flexible segments or bubbles, i.e. we simulated different *f*_bubble_ for a range of values of stiffnesses *L*_P,bubble_ and sizes *L*_bubble_ of the flexible regions. We find that for small bubbles (*L*_bubble_ ≤ 10 bp), the simulated force-extension data remain well-described by the force-extension expression of the WLC model (Figure 6C and Supplementary Figure S10, S11, and S12). In contrast, large bubble sizes (*L*_bubble_ > 25-100; Figure 6D and Supplementary Figure S10) lead to deviations from the WLC model, at odds with the experimental findings (Figure 1). Fits of the WLC to the simulated data find reduced (effective) bending persistence lengths with i) increasing bubble fraction, ii) decreasing bubble stiffness *L*_P,bubble_, and iii) decreasing bubble size *L*_bubble_ (Figure 6C,D and Supplementary Figure S11). We note that since for large bubbles the simulated DNA does not behave like a WLC, the fitted apparent *L*_P_ is difficult to interpret.

Introducing bubbles into the simulated chains in the absence of applied forces, similarly leads to a reduction of the measured 3D end-to-end distance <*R*^2^> (Figure 6C,D and Supplementary Figure S13). Again, the end-to-end distances decreases with i) increasing bubble fraction, ii) decreasing bubble stiffness *L*_P,bubble_, and iii) decreasing bubble size *L*_bubble_ (Figure 6C and Supplementary Figure S14). We find that for large bubbles sizes (*L*_bubble_ > 25-100), <*R*^2^> decreases both in absolute value (Supplementary Figure S14) and relative to the zero DMSO conditions (Figure 6D) too little to account for the experimental observations. For small bubble sizes, we observe a reduction in <*R*^2^> broadly consistent with experiments and of similar relative magnitude as the reduction in the apparent bending persistence length (Figure 6C). Therefore, both the results from force-extension simulations and from simulations of molecules in free solution suggest that only models with small and dispersed bubbles are consistent with the experimental data. This conclusion is further reinforced by rotation-extension simulations, i.e. control simulations with constrained linking number, which qualitatively align with tweezer measurements for small bubbles but show significant deviations for large bubbles (Supplementary Figure S15).

Further, we find that simulations with bubble stiffnesses > 2 nm would require bubble fractions > 10% (*f*_bubble_ > 0.1) to match the experimentally observed reductions in apparent bending stiffness and <*R*^2^>, which is likely unrealistic. In addition, we find that the dependencies of the apparent bending persistence lengths and <*R*^2^> are convex with increasing bubble fraction (Figure 6C,D and Supplementary Figure S14), in particular at large bubble fractions. Conversely, the reduction of the experimentally determined values of the bending persistence lengths and <*R*^2^> are approximately linear (or even slightly concave, Figure 1D). Together, this suggests again that likely small and flexible bubbles with a small bubble fraction are most consistent with the experimental data and/or that the dependence between DMSO concentration and bubble fraction is super-linear.

Taken together, our coarse-grained simulations show that introducing flexible segments into the WLC-model can account at least semi-quantitively for the experimentally observed reduction in apparent *L*_P_ and <*R*^2^>. The simulations find that small and disperse bubbles with a stiffness similar to *L*_P,bubble_ ∼ 1 nm give the best agreement with experimental data. For these parameters a small fraction of bubbles (∼3-4 % at 50% DMSO) is sufficient to account for the experimental observations up to 50% DMSO. A small fraction of bubbles, in turn, is consistent with the fact that we do not observe substantial unwinding of the helix from rotation-extension curves and bubbles are not observed by AFM imaging.

## CONCLUSION

We have used highly complementary single-molecule methods to probe the effect of DMSO on DNA conformations and mechanics. Magnetic tweezers force spectroscopy, where the ends of the DNA are constrained and external forces applied, and AFM imaging, where DNA is visualized after equilibration in free solution, provide an overall consistent view. The bending persistence length, mean-squared end-to-end distances, and shape parameters like areas and long-axis lengths all decrease systematically with increasing DMSO concentration. While very high DMSO concentrations (> 50% DMSO in near-physiological buffer) lead to a collapse of DNA, the observed changes are gradual and approximately linear up to 50%. For low DMSO concentrations, < 10%, the observed changes are minor, of the same order as typical measurement errors in single-molecule measurements. Our observations validate the -often implicit-assumption to neglect the effect of DMSO on DNA in single-molecule assays, where DMSO is introduced in concentrations usually below 10%, e.g. upon adding fluorescent dyes from a DMSO containing stock (8,82) and are also consistent with the observation that higher order DNA origami structures remain stable in up to 30-40% DMSO (19). Our measurements are at odds with a recent report that finds a stronger reduction of end-to-end distances and bending persistence lengths upon addition of DMSO (20). We note that the measurements of Xu et al. are performed in parts at very low ionic strength (≤ 1 mM monovalent salt), which might make the DNA particularly susceptible to the effects of DMSO, as has recently been observed for DNA origami structures (19). In contrast, our measurements are all performed under matching conditions in physiologically relevant buffer (∼150 mM monovalent salt).

Our coarse-grained Monte Carlo simulations rationalize the observed effects, at least qualitatively, in the light of a simple physical model: Introducing softer, much more bendable segments termed bubbles into the stiffer chain representing double-stranded DNA leads to a reduction of the apparent bending persistent length and mean squared end-to-end distance. Comparison of the model to our experimental observations suggests that the flexible bubbles are dispersed throughout the chain and remain relatively small. Under those condition, only a small fraction of bubbles is sufficient to account for the experimentally observed reductions in apparent *L*_P_ and <*R*^2^>. We anticipate that the trends observed in our model will also be useful to interpret other types of experimental situations where DNA is locally rendered more flexible, e.g. by increased temperature or DNA damage. In summary, our results provide quantitative parameters and a baseline understanding of how DMSO affects DNA conformations that will aid the interpretation and modeling of more complex biophysical and biochemical assays involving DMSO.

## AUTHOR CONTRIBUTION STATEMENT

K.R.S.: conceptualization, data analysis, investigation (MT measurements), visualization, writing – original draft preparation. C.K: conceptualization, data analysis, investigation (AFM measurements), visualization, writing – original draft preparation. E.S.: conceptualization, data analysis, investigation (MC simulations), writing – original draft preparation. S.D.P., P.J.K., W.V., and H.S.: data analysis, supervision, writing – review and editing. J.L.: conceptualization, funding acquisition, supervision, project administration, writing – original draft preparation.

## ACKNOWLEDGEMENTS

We thank Dave van den Heuvel, Elleke van Harten, Roy Hoitink for laboratory support and Gerhard Blab, Alptug Ulugöl, Willem Gispen, and Tor Sewring for useful discussions.

This work was supported by Utrecht University, by the European Research Council Consolidator Grant “ProForce”, a Feodor-Lynen fellowship from the Alexander von Humboldt Foundation, and the Deutsche Forschungsgemeinschaft (DFG, German Research Foundation) under Germany’s Excellence Strategy - EXC-2068 – 390729961.

