## Supplementary Information for "The Effects of DMSO on DNA Conformations and Mechanics"

**Content**

**Supplementary Table S1**

**Supplementary Figures S1 – S15**

### Supplementary Table

**Supplementary Table S1. Statistical tests comparing DNA end-to-end distance distributions under DMSO-treatment**. Results of two-sample two-tailed *t*-tests comparing the means and of two-sample Kolmogorov-Smirnov tests comparing the full distributions of end-to-end distances of DMSO treated DNA samples with the PBS only condition.

| 2-sample *t*-tests: | |
|---|---|
| PBS vs. 2% DMSO | $p = 8.3 \times 10^{-6}$ |
| PBS vs. 10% DMSO | $p = 1.7 \times 10^{-2}$ |
| PBS vs. 20% DMSO | $p = 2.4 \times 10^{-12}$ |
| PBS vs. 50% DMSO | $p = 3.7 \times 10^{-18}$ |

| Kolmogorov-Smirnov tests: | |
|---|---|
| PBS vs. 2% DMSO | $p = 7.8 \times 10^{-6}$ |
| PBS vs. 10% DMSO | $p = 1.2 \times 10^{-3}$ |
| PBS vs. 20% DMSO | $p = 1.6 \times 10^{-11}$ |
| PBS vs. 50% DMSO | $p = 7.0 \times 10^{-16}$ |

### Supplementary Figures

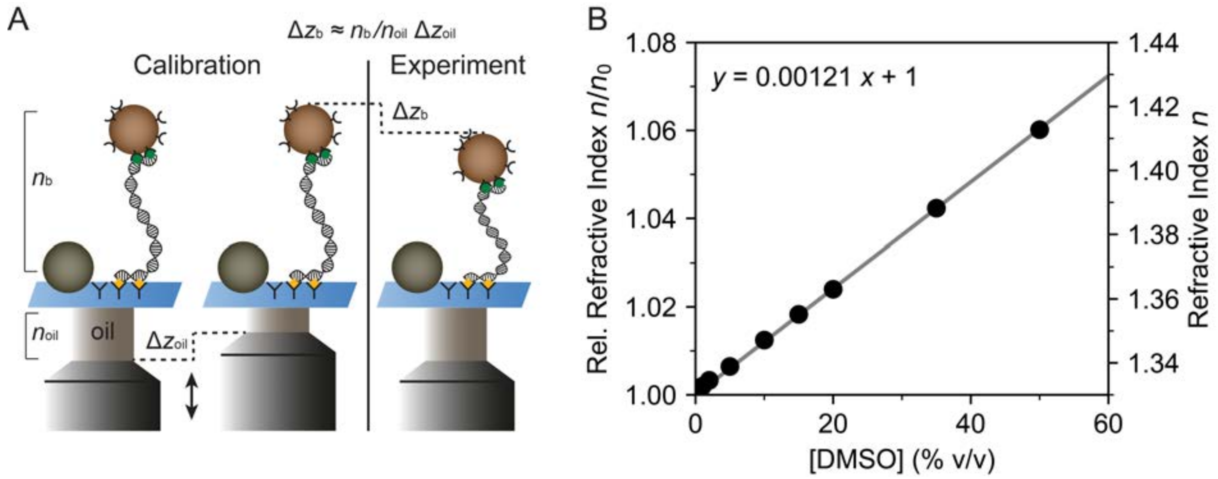


**Supplementary Figure S1. Effect of refractive indices on magnetic tweezers measurements.** A) Schematic of the determination of vertical displacement using magnetic tweezers. Prior to the experiment, a calibration routine is performed in which diffraction patterns are collected while displacing the oil immersion objective vertically, i.e., a displacement of $\Delta z_{\mathrm{oil}}$, to construct a so-called look-up table (LUT). During the experiment, the vertical position of the bead is determined by correlation of the diffraction pattern with the previously recorded LUT. To obtain the change in the vertical position of the bead $\Delta z_{\mathrm{b}}$ from the changes determined by comparison with the LUT, one needs to correct for the difference in index of refraction between oil and buffer. The correction factor is, at least approximately, given by the ratio of the refractive index of the oil $n_{\mathrm{oil}}$ = 1.5154 and of the buffer $n_{\mathrm{b}}$ and we, therefore, determine the absolute $\Delta z_{\mathrm{b}}$ according to $\Delta z_{\mathrm{b}} \approx n_{\mathrm{b}}/n_{\mathrm{oil}} \cdot \Delta z_{\mathrm{oil}}$. B) Refractive index of DMSO solutions in PBS. The solid line is a linear fit to the data relative to the refractive index of water (coefficient of determination $R^2$ = 0.999). For measurements in varying concentrations of DMSO, we still employed a LUT measured using PBS only buffer and we take into account the index of refraction of the (DMSO containing) buffer by using the measured values for different DMSO concentrations for $n_{\mathrm{b}}$.

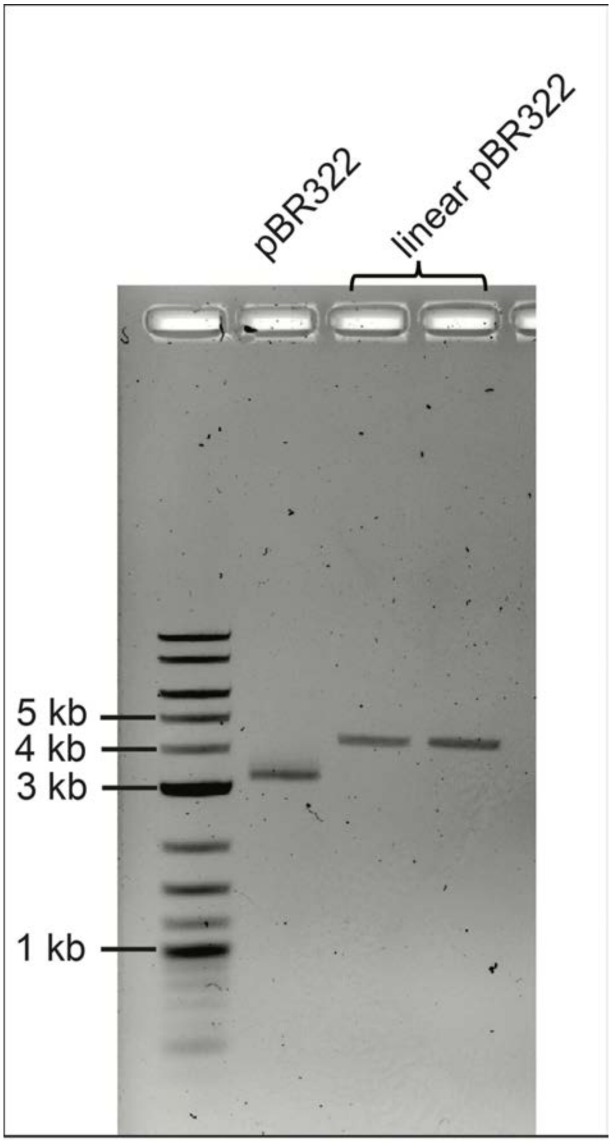


**Supplementary Figure S2. Agarose gel image of pBR322 plasmid DNA** (1st lane) and two independent linearization batches of pBR322 (2nd and 3rd lane) after linearization with EcoRI at one restriction site in the plasmid. DNA is stained with ROTI-Gel stain and visualized under UV light. 1 kb Plus DNA Ladder (NEB) is used as a size reference, and relevant bands are annotated. Due to the supercoiled structure, the plasmid DNA travels faster through the gel than the linearized DNA. The two independent linearization batches show excellent reproducibility, travel at the expected DNA length (4361 bp for pBR322) compared to the DNA ladder, and run as a single band, suggesting a successful and complete digest.

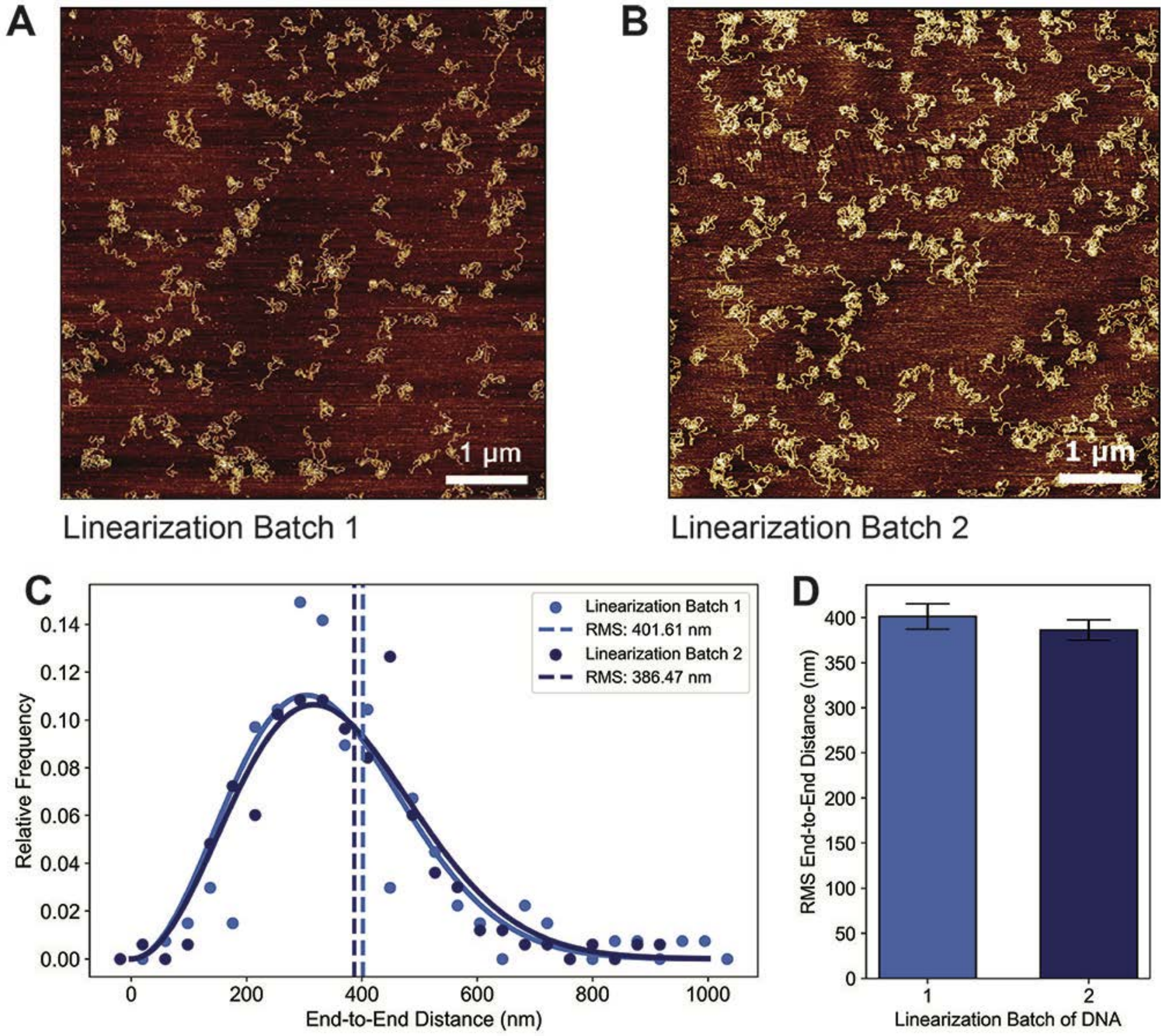


**Supplementary Figure S3. Comparison of two independent linearization batches of pBR322.** A, B) AMF images of linear DNA (4361 bp) incubated in PBS buffer and imaged on PLL-mica in air. The two DNA batches were prepared in independent linearization reactions (Materials and Methods) and show similar behavior when deposited on the surface. C) Histograms of DNA end-to-end distances in PBS show excellent agreement of distributions for two independent linearization batches. RMS values are visualized as dashed vertical lines. Solid lines are fits of the Gaussian chain model (Equation 1). D) Bar plot comparing the RMS end-to-end distances in PBS for two independent linearization batches of DNA. There is no statistically significant difference between the two batches, neither with regards to the RMS end-to-end distances (mean ± standard deviation) of 402 nm ± 14 nm and 386 nm ± 11 nm, respectively, (the difference was assessed with a two-sample two-tailed $t$-test, $p = 0.6$), nor with regards to the overall distributions (as assessed by a two-sample Kolmogorov-Smirnov test, $p = 0.8$).

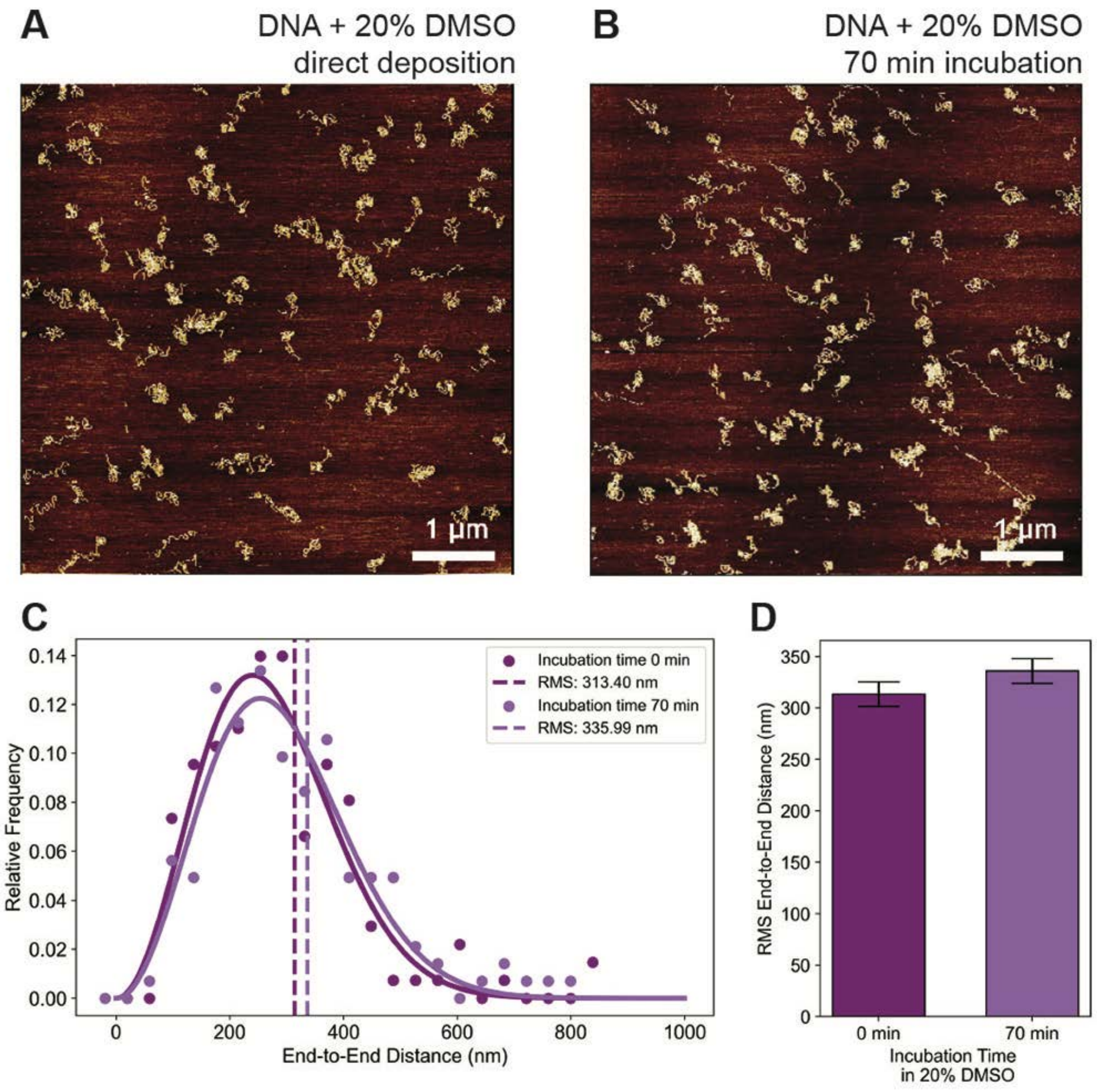


**Supplementary Figure S4. Comparison of different incubation times of linear DNA with 20% DMSO before surface deposition.** A, B) AMF images of linear DNA (4361 bp) in the 20% (v/v) DMSO condition on PLL-mica measured in air. The two batches of DNA were prepared either by incubating briefly (~30 s, A) or for a much longer time (70 min, B) in the final DMSO concentration prior surface deposition. C) Histograms of DNA end-to-end distances in 20% DMSO with the short (dark purple) and long (light purple) incubation time. The RMS values of 313 ± 12 nm and 336 ± 12 nm are visualized as dashed vertical lines. Solid lines are fits of the Gaussian chain model (Equation 1). D) Bar plot comparing the RMS end-to-end distances in 20% DMSO with and without incubation time before depositing the sample. There is no statistically significant difference between the batches with the short and long incubation time, neither with regards to the RMS end-to-end distances (as assessed with a two-sample two-tailed $t$-test, $p = 0.2$) nor with regards to the overall distributions (as assessed by a two-sample Kolmogorov-Smirnov test, $p = 0.3$).

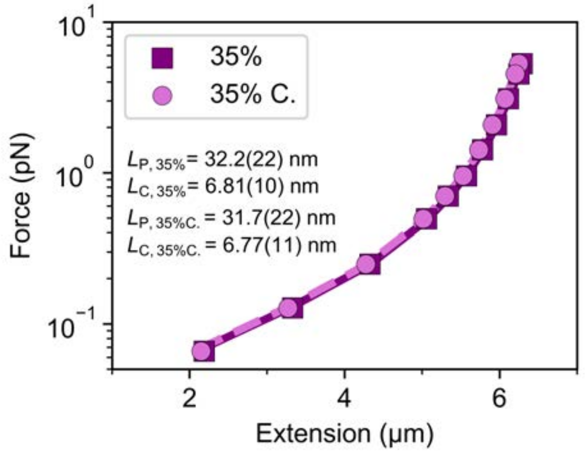


**Supplementary Figure S5. Control measurements adjusting the ionic strength after addition of DMSO.** Force-extension curves for 20.6 kbp DNA, similar to the data shown in Figure 1, for a DMSO concentration of 35%. For the data labeled "35%", 35% DMSO were added to 1x PBS buffer, as was done for the other DMSO concentrations used in the main text of the paper. For the control data labeled "35% C.", the initial PBS concentration was increased, such that it corresponds to 1x PBS after addition of 35% DMSO. Both samples give essentially identical force-extension curves and the fitted values of the contour and bending persistence lengths are within experimental error (the values in the parentheses indicate the standard deviation of 11 independent molecules as the last significant digit).

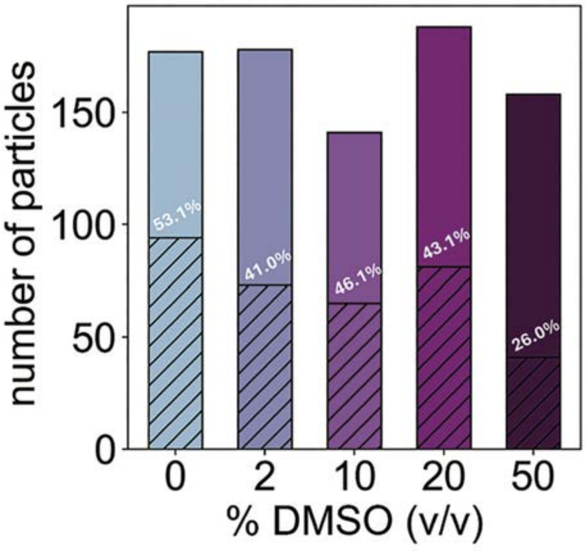


**Supplementary Figure S6. Number of molecules analyzed for manual shape analysis and fraction of molecules for end-to-end distance analysis in varying DMSO concentrations.** Colored boxes (blue to purple with increasing DMSO) indicate the number of single molecules used for manual shape analysis, the striped boxes indicate the number of the single molecules with visible free ends for end-to-end distance analysis. The data shown are from analysis of all single molecules from 3 images per condition; the end-to-end distance data in Figure 5 uses additional images with similar percentages. We can measure the end-to-end distances for approximately half of the single molecules but are limited to ~25% in 50% DMSO where we observe a more substantial compaction of DNA molecules.

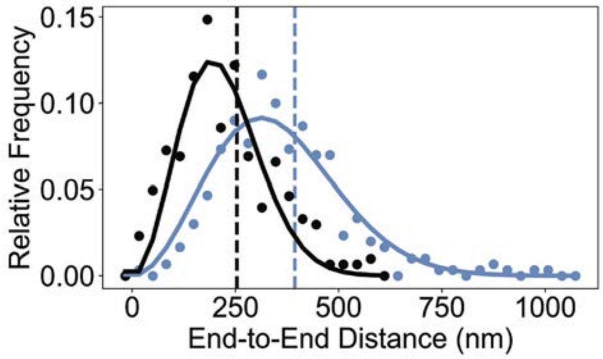


**Supplementary Figure S7. Comparison of DNA end-to-end distance distribution for linearized pBR322 plasmid DNA in PBS buffer and in buffer containing 5 mM $Mg^{2+}$**. Blue data points are the data in PBS buffer, from this work (Materials and Methods). The black data points are the data from Brouns *et al*. (ACS Nano, 12(12):11907-11916, 2018; Ref. 49 of the main text). Both data sets were obtained by AFM imaging in air of linearized DNA (4361 bp) deposited on a PLL-coated mica surfaces. Overall, the distributions of end-to-end distances look similar. However, the data in the 5 mM $Mg^{2+}$ conditions show smaller mean end-to-end distances (253 ± 7 nm, black vertical dashed line) in comparison with our results (393 ± 8 nm, blue vertical dashed line) , which is to be expected since $Mg^{2+}$ reduces the bending persistence length of DNA and tends to compact DNA. The lines are fits of the Gaussian chain model (Equation 1) to the data.

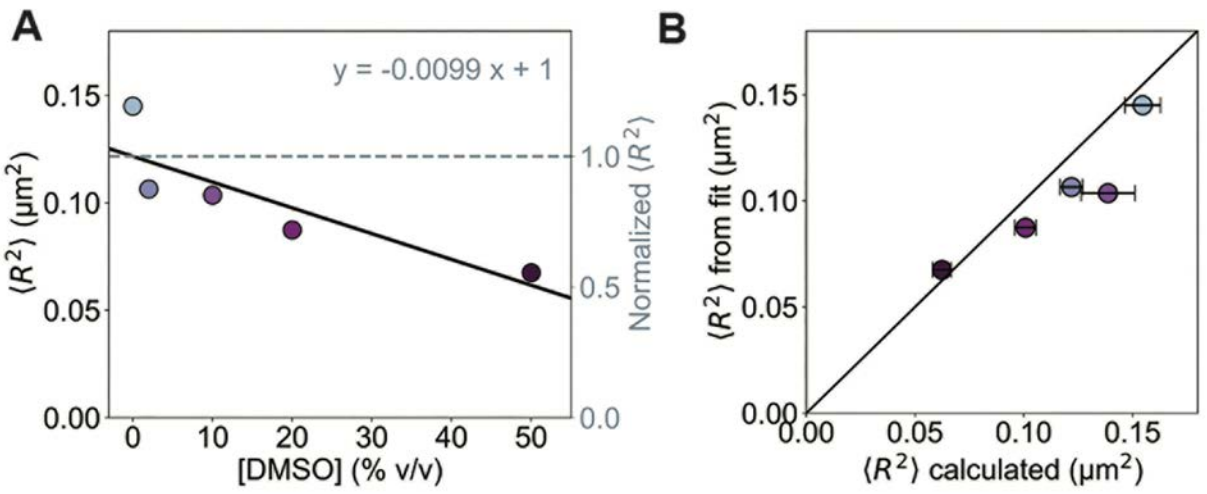


**Supplementary Figure S8. Comparison of calculated and fitted <$R^2$>** . A) <$R^2$> values obtained from fits of the Gaussian chain model (Equation 1 in Materials and Methods) as a function of DMSO concentration. The straight line indicates a linear fit corresponding to ~1% decrease in <$R^2$> per %-DMSO. This figure is similar to Figure 5C, where we show <$R^2$> values directly calculated from the end-to-end distances measured in AFM images (and find a change of 1.15% per %-DMSO). B) Correlation of <$R^2$> values from the direct calculation and from the Gaussian model fits. Overall, the two approaches to obtaining <$R^2$> are in close agreement. The fitted values show a tendency to smaller values compared to the directly computed <$R^2$> values. The black line is a 45-degree line as a guide to the eye.

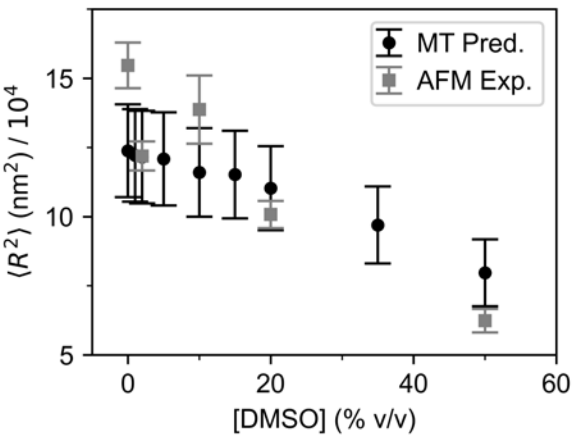


**Supplementary Figure S9. Comparison of measured DNA end-to-end distances by AFM imaging and predictions based on parameters from MT force spectroscopy.** Grey squares are the $<R^2>$ values measured experimentally by AFM imaging in air after deposition in PLL-mica (Materials and Methods; Figures 3 and 4). Black circles are predictions of the 3D mean squared end-to-end distance from Equation 1 using the values for the bending persistence length $L_P$ and contour length $L_C$ parameters determined by MT force spectroscopy (Figure 1). Error bars for the black data points are from the propagation of experimental uncertainties of the measured parameters.

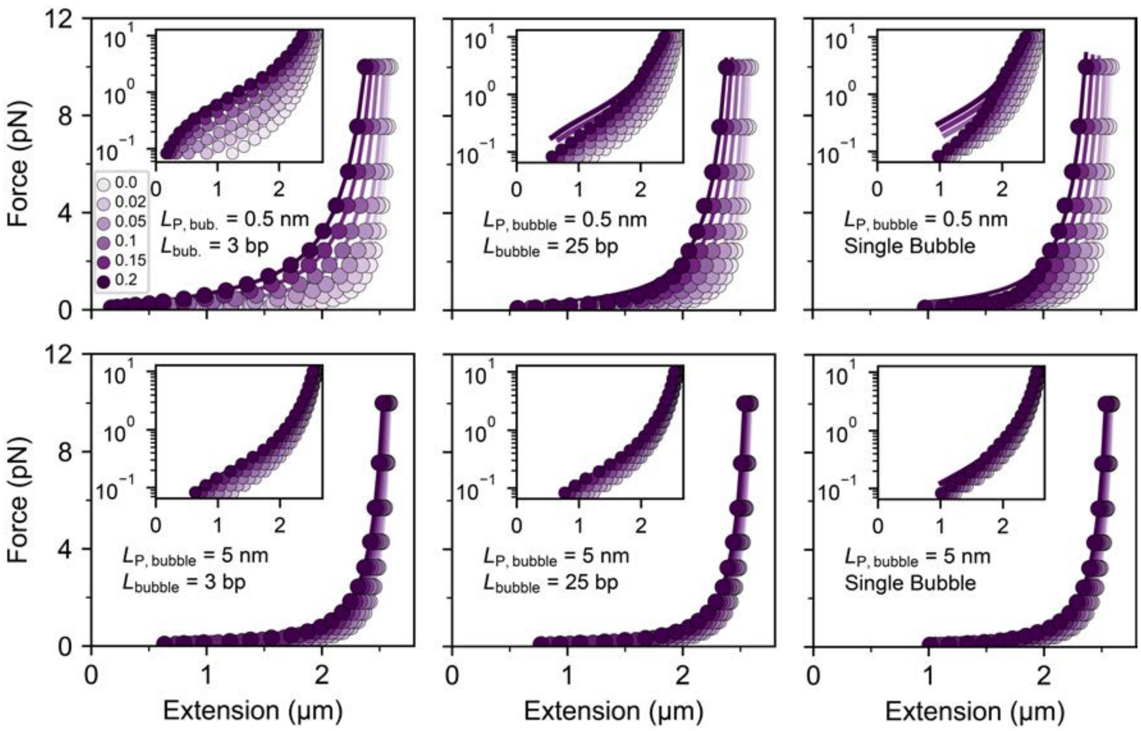


**Supplementary Figure S10. Force-extension relationships for coarse-grained DNA simulations with bubbles.** Force-extension relationships from coarse-grained Monte Carlo simulations (symbols) and fits of the WLC model (solid lines). The different colors in each panel indicate different bubble fractions $f_{\text{bubble}}$ (increasing from light to dark shades; see legend in the top left panel; the same color code applies to all panels). The simulations systematically varied the flexibility of the bubble segments $L_{\text{P,bubble}}$ in the range 0.5 nm to 5 nm. Shown here are the data for $L_{\text{P,bubble}}$ = 0.5 nm (top row) and 5 nm (bottom row). In addition, we systematically varied the size of the flexible bubble regions, $L_{\text{bubble}}$, in the range of 3 bp to 10,000 bp. The 10,000 bp case corresponds to the situation that all bubble segments are in one cluster, i.e. that there is a single bubble that comprises all flexible segments. Shown are the data for $L_{\text{bubble}}$ = 3 bp (left panels), 25 bp (middle panels), and the single bubble case (right panels). The insets show the same data in log-linear representation.

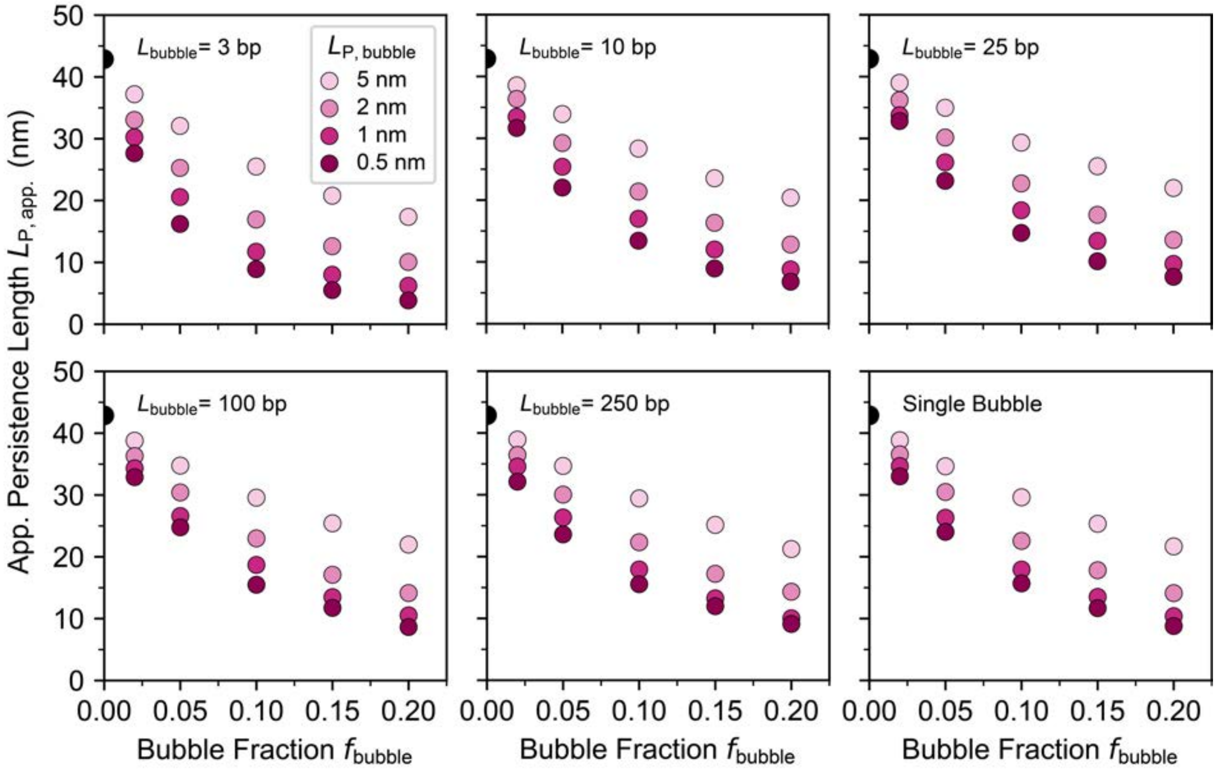


**Supplementary Figure S11. Apparent bending persistence lengths from coarse-grained DNA simulations with bubbles.** Values for the apparent bending persistence length obtained from WLC fits to the simulated force-extension responses of DNA with flexible bubbles (examples of the data and fits are shown in Supplementary Figure S10). Values for the fitted apparent $L_P$ are shown as a function of the bubble fraction and stiffness of the bubbles $L_{P,bubble}$ (color coded from dark to light shades; see legend in the top left panel; the same color code is used for all panels). The different panels show data for different bubble sizes $L_{bubble}$. The "single bubble" condition has $L_{bubble}$ = 10,000 bp, such that all bubble segments are in one cluster. The black dots at a bubble fraction of 0 corresponds to the value determined from coarse-grained simulations of the discrete WLC without flexible bubbles, which accurately recover the input bending persistent length of 43 nm.

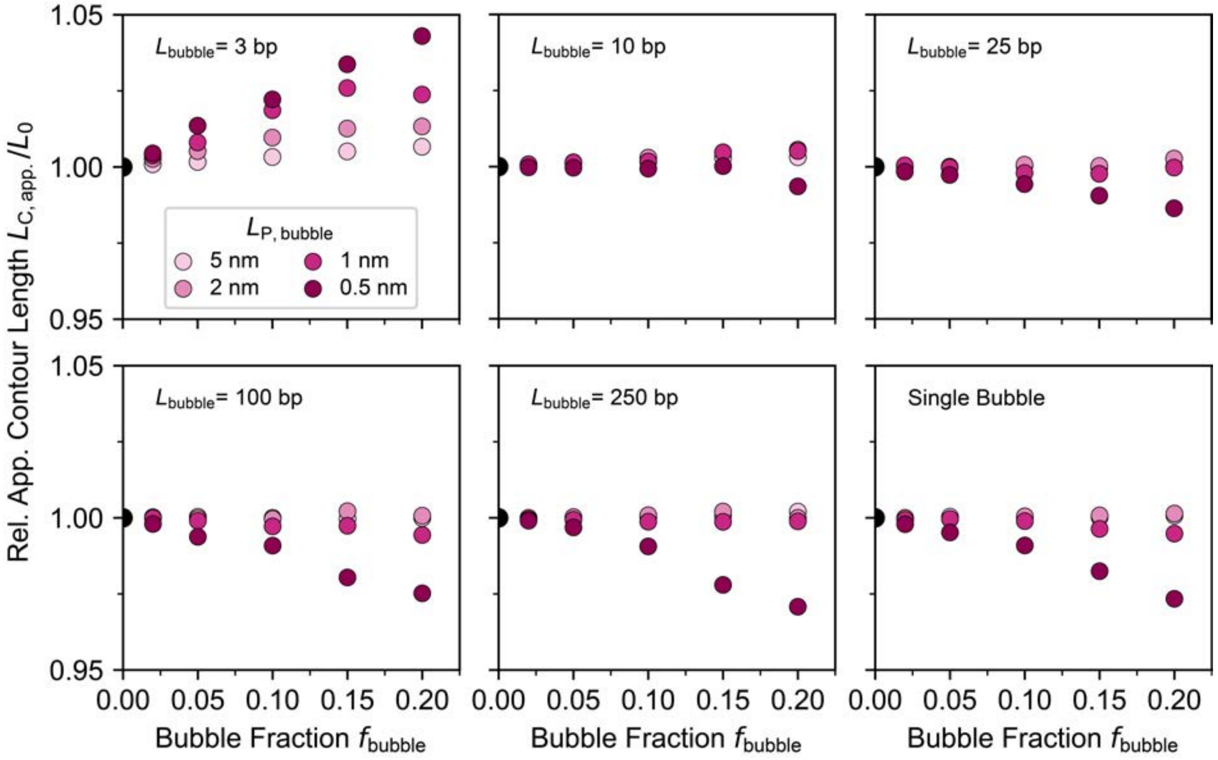


**Supplementary Figure S12. Fitted contour lengths from coarse-grained DNA simulations with bubbles.** Values for the fitted contour lengths obtained from WLC fits to the simulated force-extension responses of DNA with flexible bubbles (examples of the data and fits are shown in Supplementary Figure S10), normalized to the contour length of the chain in the absence of bubbles. Normalized values for the fitted apparent $L_C$ are shown as a function of the bubble fraction and stiffness of the bubbles $L_{P,bubble}$ (color coded from dark to light shades; see legend in the top left panel; the same color code is used for all panels). The different panels show data for different bubble sizes $L_{bubble}$. The "Single Bubble" condition has $L_{bubble}$ = 10,000 bp, such that all flexible bubble segments are in one cluster. Note the narrow scale of the y-axis. The fitted contour lengths are almost constant across the different conditions.

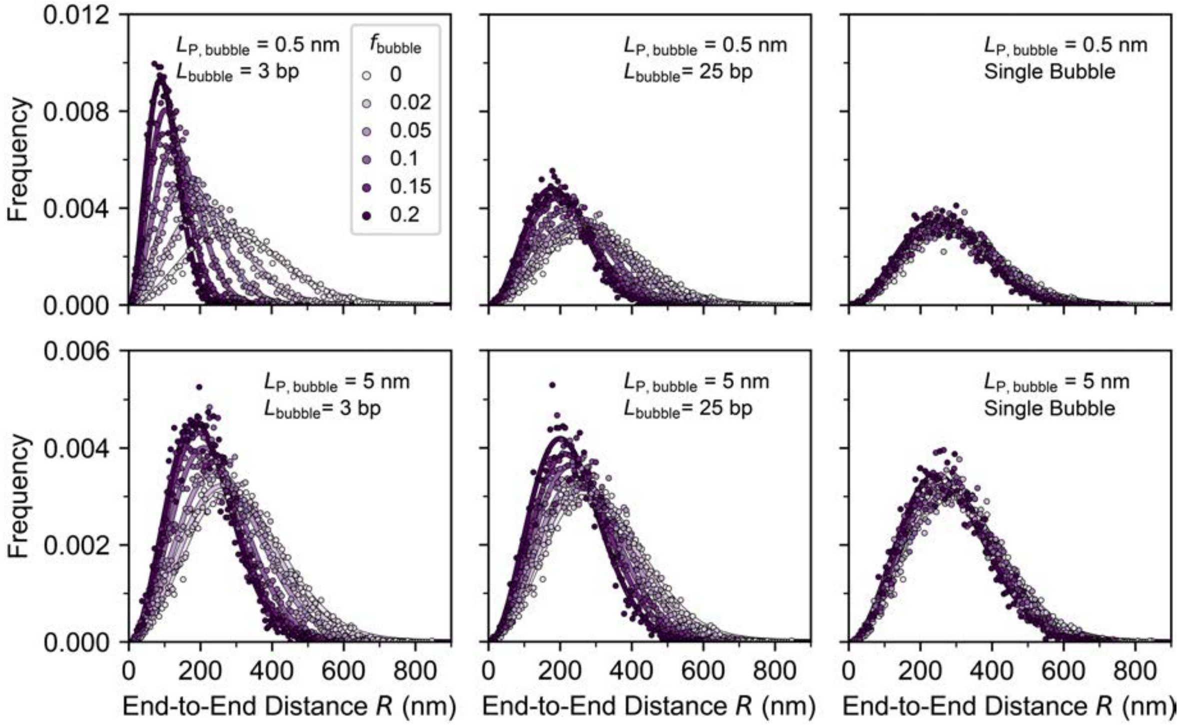


**Supplementary Figure S13. End-to-end distance distributions for coarse-grained DNA simulations with bubbles.** Distributions of end-to-end distances from coarse-grained Monte Carlo simulations (symbols) and fits of the Gaussian chain model (Equation 1; solid lines). The different colors in each panel indicate different bubble fractions $f_{bubble}$ (increasing from light to dark shades; see legend in the top left panel; the same color code applies to all panels). The simulations systematically varied the flexibility of the bubble segments $L_{P,bubble}$ in the range 0.5 nm to 5 nm. Shown here are the data for $L_{P,bubble}$ = 0.5 nm (top row) and 5 nm (bottom row). In addition, we systematically varied the size of the flexible bubble regions, $L_{bubble}$, in the range of 3 bp to 10,000 bp. The 10,000 bp case corresponds to the situation that all bubble segments are in one cluster, i.e. that there is a single bubble that comprises all flexible segments. Shown are the data for $L_{bubble}$ = 3 bp (left panels), 25 bp (middle panels), and the single bubble case (right panels).

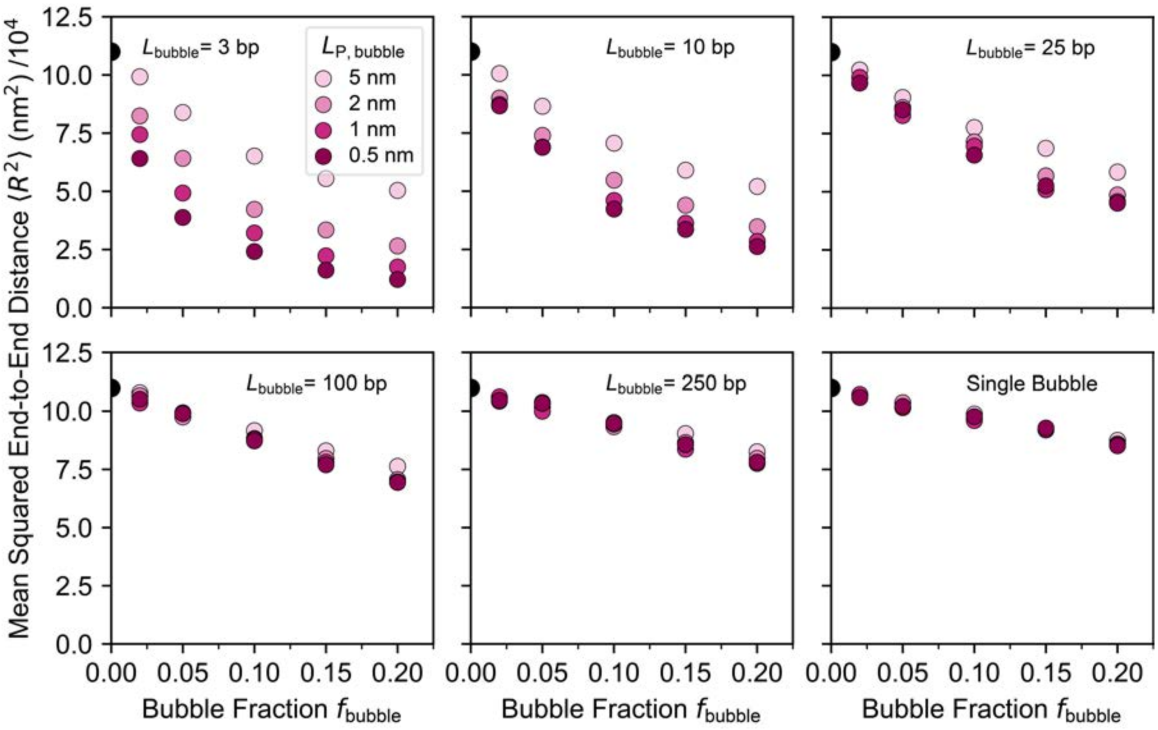


**Supplementary Figure S14. Mean-squared end-to-end distance from coarse-grained DNA simulations with bubbles.** Values for the mean-squared end-to-end distance $<R^2>$ obtained from the Monte Carlo simulations of DNA with flexible bubbles (examples of the distributions are shown in Supplementary Figure S13). Values for $<R^2>$ are shown as a function of the bubble fraction and stiffness of the bubbles $L_{P,bubble}$ (color coded from dark to light shades; see legend in the top left panel; the same color code is used for all panels). The different panels show data for different bubble sizes $L_{bubble}$. The "Single Bubble" condition has $L_{bubble}$ = 10,000 bp, such that all flexible bubble segments are in one cluster.

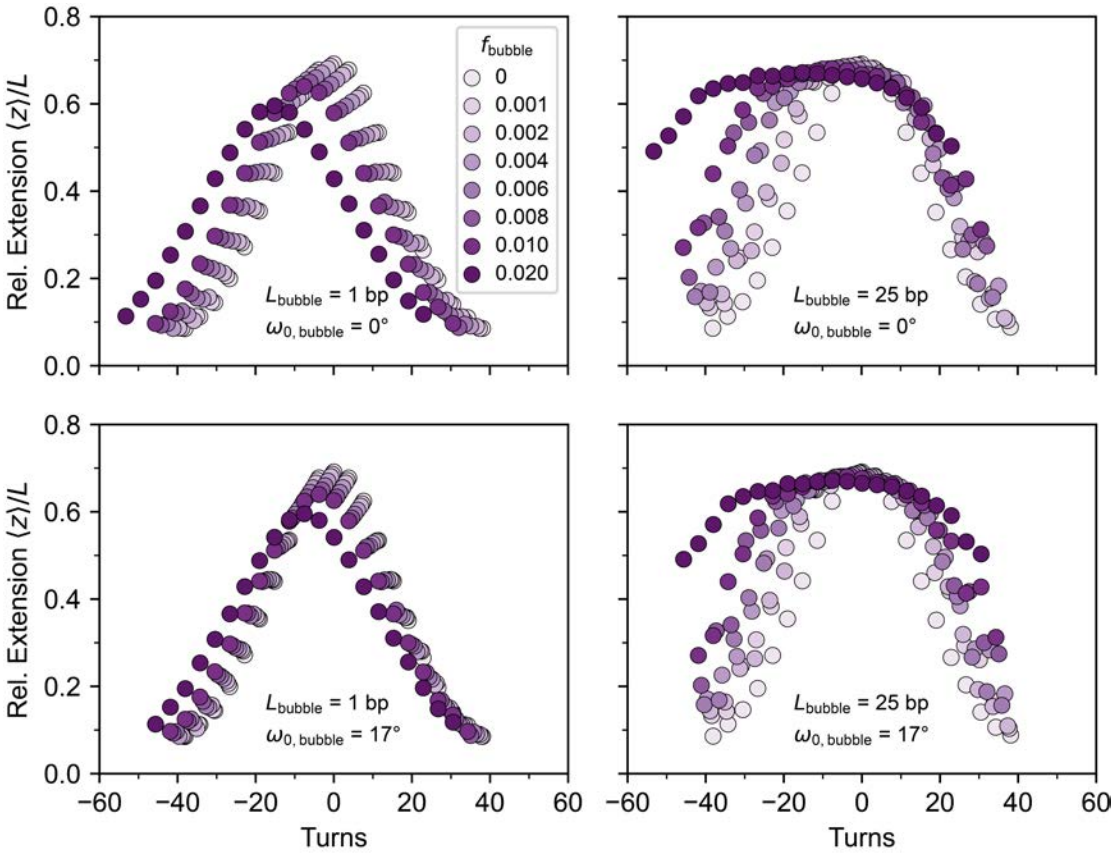


**Supplementary Figure S15. Monte Carlo simulations of rotation-extension curves for DNA molecules with bubbles of different sizes.** Rotation-extension simulations were performed for 8 kbp molecules subjected to a stretching force of $F = 0.25$ pN. Two bubble sizes were considered: 1 bp (left column) and 25 bp (right column). Additionally, two scenarios for the intrinsic twist in bubble steps were examined: one in which intrinsic twist is completely removed (top row) and one in which it is halved (bottom row). These correspond to a twist change of −34° and −17° per bubble step, respectively. Extensions are normalized to the contour length of the molecule. For small, dispersed bubbles, the rotation-extension curves remain largely unchanged, particularly at low bubble fractions, consistent with experimental observations. In contrast, large bubbles lead to fundamentally different behavior. The pre-buckling regime (the upper part of the curve, preceding the region of linear extension reduction) is significantly extended even at very low bubble fractions $f_{bubble}$. This effect arises because extended flexible regions (large bubbles) can efficiently absorb linking strain through local helicoidal or superhelical writhing. Consequently, these regions act as reservoirs for linking strain, which must be exhausted before the buckling transition of the remaining duplex DNA is initiated. These simulations further reinforce our conclusion that the modulation of DNA elasticity due to DMSO interactions is best explained by small, dispersed bubbles.
